# Degradation of PET plastic with engineered environmental bacteria

**DOI:** 10.1101/2024.09.24.614569

**Authors:** Alice M. Banks, Umar Abdulmutalib, Christian Sonnendecker, Juhyun Kim, Charlotte Bosomworth, Stuart Brown, Ren Wei, Carolina Álvarez-Ortega, Pablo Pomposiello, Claudio Avignone-Rossa, Gerald Larrouy-Maumus, Wolfgang Zimmermann, José I. Jiménez

## Abstract

Polyethylene terephthalate (PET) is one of the most widely used plastic materials in the food and textile industry. Consequently, post-consumer PET waste is a common environmental pollutant that leaks into the environment in the form of macro and microplastics with concerning health impacts. There is a pressing need to identify novel and sustainable solutions to process the abundance of PET waste contributing to this pollution. While there is extensive research into enzymes able to hydrolyse PET *in vitro*, a similar solution for discarded or difficult-to-collect PET based on whole-cell microbial catalysts is missing. In this work we report the engineering of environmental bacteria to use PET as a growth substrate. This was achieved by isolating a strain of *Pseudomonas umsongensis* able to use the PET monomer terephthalate as carbon source, engineering the strain to effectively secrete the high-activity PET hydrolase PHL7 through the addition of a recombinant TAT secretion leader sequence, and enhancing the bioavailability of PET by transforming it into an amorphous and macroporous structure by pre-treatment with an organic solvent. Our findings demonstrate the direct microbial consumption of PET, which could lead to improved and more sustainable upcycling strategies for this plastic.

## Introduction

Commonly used plastics are non-biodegradable materials that accumulate in the environment and pose a health hazard in almost every ecosystem on the planet (1). The plastic PET, in particular, is a versatile material used for multiple applications, from containers for the food and beverage industry, to the production of textiles (2). Compared to other plastics, PET has high collection rates and can be recycled by mechanical and chemical methods (3), however, due to its widespread use and the fact that some PET forms such as fibres released from fabrics are not easily collected, PET can find its way into the environment (4,5) which poses a health hazard (6).

Biological recycling of PET that has reached its end of life and cannot be recycled mechanically has been proposed as a mitigation strategy for its unwanted release into the environment, while contributing to the conservation of fossil resources (7). This involves the use of enzymes that constitute promising catalysts for the breakdown of polyester plastics that operate under very mild reaction conditions compared to chemical processes. Out of the most commonly used plastics, only PET is currently being used in pilot industrial enzymatic recycling processes (8). This process consists of the *in vitro* enzymatic hydrolysis of PET resulting in the production of large quantities of the PET constituent monomers terephthalate (TA) and ethylene glycol (EG) (9–11). While it is possible to repolymerise these two molecules in order to generate virgin PET, they can also be used as source of carbon or energy for microbial growth (12) as well as for conversion into molecules with added value (13–16). This two-step process, *in vitro* hydrolysis and *in vivo* upcycling (see for example (17)), requires the initial expression and purification of appropriate enzymes which increases the energy and carbon footprint as well as decreases the potential economic benefit of the valorisation of PET waste by biological means (18).

In this paper we describe the *in vivo*, or direct hydrolysis, of PET (Fig. 1A). This process has been proposed as a potential way of improving both the sustainability and economic margins of PET as well as a potential solution to reduce microplastic pollution in waterways and farming land if used during wastewater treatments (19). The ability to grow mainly on PET as carbon source with little supplementation by additional nutrients, a process also referred to as ‘PETtrophy’ (20), is exemplified by the bacterium *Ideonella sakaiensis* isolated from a plastic recycling plant (11). *I. sakaiensis* is not, however, an organism easy to culture or manipulate in the lab due, among other factors, to limited secretion capabilities (21), and previous works have advocated for the modification of workhorses for biotechnology such as *Pseudomonas putida* for PET assimilation although this goal has not been achieved to date (20). The methodology requires microorganisms capable of secreting PET hydrolases to be cultured in the presence of the plastic substrate. Moreover, the organisms could also be engineered for the assimilation of PET monomers and the generation of products of interest, which could have a significant impact on the economic recovery of PET upcycling operations. We succeeded in engineering an environmental isolate for PET assimilation following these steps: i) the screening and isolation of bacteria able to use TA as a sole carbon source; ii) the generation of broad-host plasmids for the expression and efficient secretion of PET hydrolases in the environmental isolates and iii) the pre-treatment of PET to obtain a substrate with increased bioavailability. Our results show that the systematic investigation of these key processes can lead to the overall optimisation of the process of PET biodegradation resulting in a strain able to use PET as a growth substrate.

**Fig. 1.**
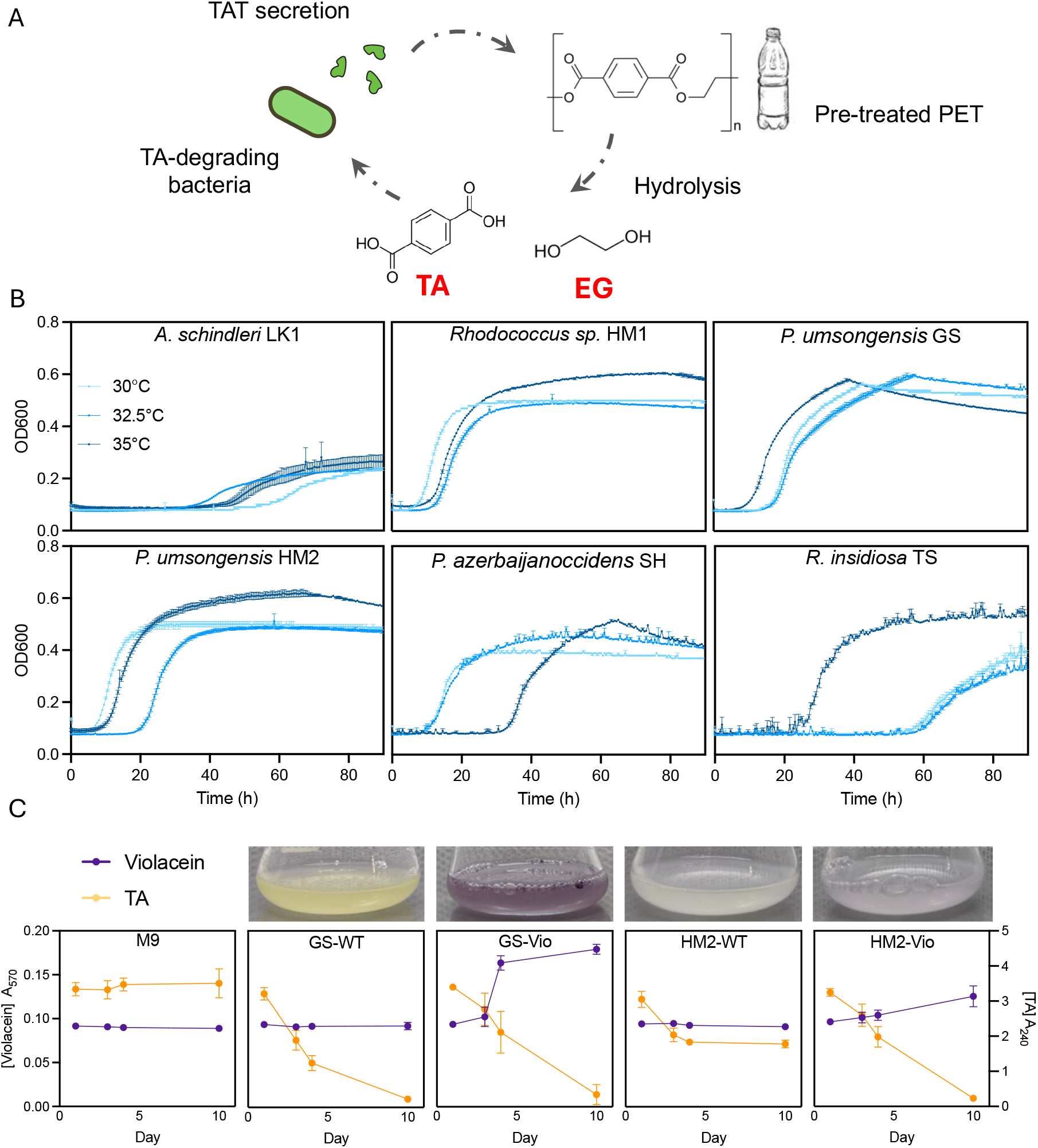
PET valorisation by environmental isolates. (A) Diagram outlining the general strategy followed in this work. TA and EG stand for, respectively, terephthalic acid (or terephthalate) and ethylene glycol. (B) Growth of the environmental isolates using the PET monomer TA as the sole carbon source across a temperature range. (C) The potential of the two *P. umsongensis* strains GS and HM2 for PET upcycling was assessed by transforming them with a plasmid containing the violacein biosynthetic pathway and monitoring the conversion of TA (yellow line) into violacein (purple line) over time. Error bars denote the standard deviation (SD) of three biological replicates.

## Results

### Isolation and growth characterisation of candidate microbial strains

Microbial strains were isolated from environmental samples taken from grassland, soil, and lake locations using TA for culture enrichment. As a result of the screening, we isolated six new bacterial strains able to grow on TA as the sole carbon source in pure cultures. These strains were sequenced completely and classified according to their 16S rRNA sequence as *Acinetobacter schindleri* LK1, *Rhodococcus* sp. HM1, *Pseudomonas umsongensis* GS, *Pseudomonas umsongensis* HM2, *Pseudomonas azerbaijanoccidens* SH and *Ralstonia insidiosa* TS. In addition to TA, we also tested their ability to use EG as growth substrate (Fig. 1B and S1).

Isolates LK1, HM1, GS, HM2, SH and TS all exhibited growth using TA as a sole carbon source, however HM1 was the only isolate showing growth on EG. The *Pseudomonas* spp. (GS, HM2, SH) and *Rhodococcus* sp. HM1 all reached a similar biomass using TA, with little difference in growth rate under the conditions tested. *R. insidiosa* TS achieved a moderate biomass using TA as a sole carbon source and also exhibited a longer lag phase than the *Pseudomonas* spp.

We conducted a genomic search of the genomes involved in TA metabolism in the environmental isolates. To this end, we investigated the presence of homologs for the *tphA1A2A3B* genes responsible for the conversion of TA into protocatechuate (PCA) (22) using a blastp search. This analysis revealed that isolates HM1, HM2, GS, and SH all contained gene clusters homologous to the *tph* operon (Table S1). The equivalent cluster was lacking in isolates LK1 and TS. At this stage we discarded the gram-positive HM1 and LK1, for which no molecular tools for genetic manipulation were available. The remaining four environmental isolates were also screened for resistance to a panel of routinely used antibiotics (Fig. S2). *R. insidiosa* TS demonstrated resistance to all antibiotics tested apart from tetracycline, whereas isolate *A. schindleri* LK1 was found to be susceptible to the full range tested. *Pseudomonas* spp. (GS, HM2, SH) only showed resistance to ampicillin which is a common feature of the genus (23).

Based on initial growth characterisation and genome analysis, the isolates *P. umsongensis* GS and *P. umsongensis* HM2. were selected for further investigation due to their ability to grow on TA as a sole carbon source; their susceptibility to a wide range of antibiotics; and the possession of the *tph* gene cluster in the genome. Furthermore, these isolates are all *Pseudomonas* spp., a genus for which an extensive molecular toolkit is available.

In a next step we tested the ability of the isolates *P. umsongensis* GS and HM2 for the production of molecules of biotechnological interest using TA as a growth substrate. The plasmid pSEVA63-Hvio, carrying the violacein biosynthetic pathway (24), was delivered to both strains, which were cultured in M9+TA and monitored over a 10-day period for the conversion of TA to violacein. The engineered strains demonstrated the conversion of TA to violacein. Wild-type cultures showed the depletion of TA over the 10-day incubation period, but no accumulation of violacein. Strains carrying the pSEVA63-Hvio plasmid similarly demonstrated depletion of TA, but also caused the accumulation of the violacein pigment (detectable at 570 nm), and the cultures developed a visible purple colour resulting from production (Fig. 1C). Out of the two isolates tested, *P. umsongensis* GS showed the highest violacein production ability when expressing the recombinant pathway, which suggests that it could also be a good host for the expression of PET hydrolases.

### Engineering environmental isolates for the expression and secretion of PET hydrolases

There are a considerable number of enzymes active against PET reported in the literature (25). We focused on two main groups for this work to assess whether they were capable of supporting microbial growth on PET (Table S2). The first group comprises enzymes derived from PETase and MHETase of *Ideonella sakaiensis*, a mesophilic proteobacterium, including variants engineered through directed evolution (26–28). In contrast, the second group consists of enzymes sourced from thermophilic actinobacteria or identified through metagenomic screenings (9,29–31).

The enzymes studied in this work were originally used for cytoplasmic expression in *Escherichia coli* devoid of a signal peptide, which we introduced in their N-terminus of each protein for their secretion in the environmental strains. Since all PET hydrolases tested have been expressed successfully in the cytoplasm of *E. coli* (9,26–31), we therefore sought to take advantage of the type II secretion system of *Pseudomonas* responsible for the extracellular secretion of folded proteins. To this end, we added the N-terminal signal peptide of the UxpB phosphatase of *Pseudomonas putida* KT2440 encoded by PP_1043 (32) to each of the enzymes. In its native organism, the UxpB phosphatase is secreted extracellularly under low phosphate conditions. UxpB is transported to the cell surface by the type II Xcp secretion system thanks to the presence of a 52-amino acid long signal peptide carrying a twin-arginine translocation motif (TAT).

We tested the secretion of the enzymes in different gram-negative species including *E. coli* (used for cloning) and *P. putida* (native organism of the signal peptide), as well as the environmental isolate *P. umsongensis* GS. The secretion of all the thermophilic enzymes (TfCut2, TCur0390, LCC and PHL7) was easily detected by the presence of clearance halos around producing colonies when the cells were cultured on plates containing the model substrate polycaprolactone (PCL) but not in the equivalent colonies expressing IsPETase and their variants (Fig. 2A). Interestingly, the chimeric protein generated by the fusion of the MHETase and PETase of *I. sakaiensis* (33) was also active against PCL although we failed to detect the presence of this depolymerisation activity in the corresponding individual enzymes. We also tested the activity of the different enzymes against PET nanoparticles. Clearance halos were detected only in colonies producing thermophilic enzymes after incubation at 50°C (Fig. 2B), but not in the case of mesophilic enzymes (not shown) or in a negative control of TfCut2 (TfCut2mut) generated in this work carrying the inactivating mutation S187A.

**Figure 2.**
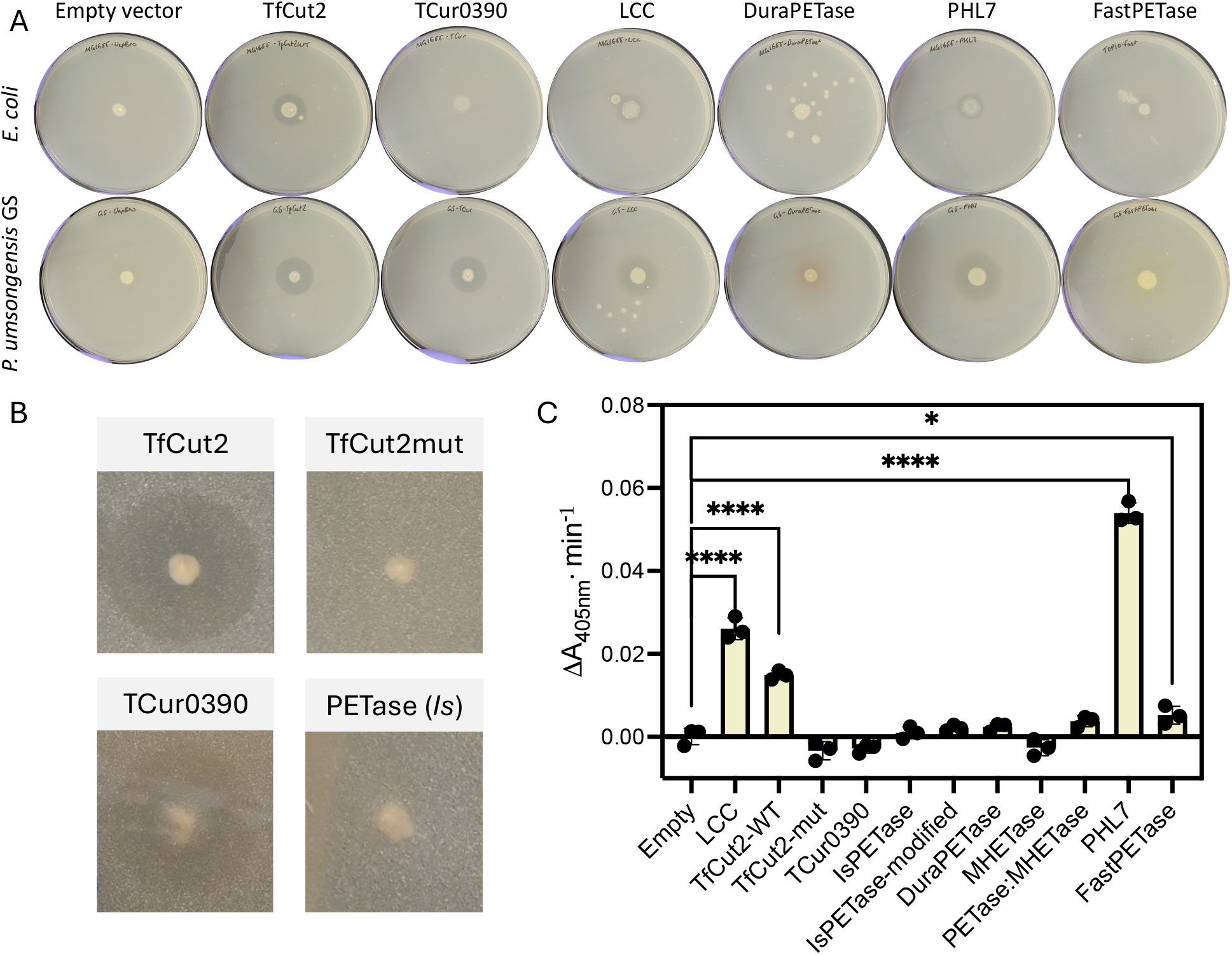
Activity of PET hydrolases recombinantly expressed and secreted. (A) PCL degradation assay on plates. (B) PET nanoparticle activity of selected hydrolysing enzymes when expressed in *P. umsongensis* GS. (C) Activity against *p*NPB at 30°C of *E. coli* supernatants producing the different enzymes. IsPETase modified refers to the variant W159H-S238F (26). Error bars indicate the standard deviation of three independent replicates. Within group differences were analysed with an ANOVA followed by a post-hoc Dunnett test. For significance please check the methods section.

In a following experiment, we set to identify the best enzyme to support microbial growth, which is expected to correlate with the total level of activity found in culture supernatants of the bacteria. It is worth noting that levels of enzymatic activity using purified enzymes vary widely between publications as there is no standardised way of measuring polyester depolymerisation activity since conditions change from one study to another. To mitigate this problem, we conducted a side-by-side comparison of enzymatic activity of enzymes secreted to the culture broth by *E. coli* containing the different plasmids in a 48 h culture. In these experiments, the activity determined results from the combination of several factors including the kinetic properties of the enzyme in culture medium and the ability of the strain to secrete a particular enzyme, all of which can vary for each enzyme, and none of these properties were determined individually.

Under the conditions tested, PHL7 showed the greatest promise to support microbial growth on PET. Given the general lack of activity of IsPETase derivatives against PCL, we used the model substrate *p*-nitrophenyl butyrate (*p*NPB) in order to detect the presence of esterolytic activity of all of the enzymes in culture supernatants at different temperatures (Fig. S3). In this assay, the presence of activity was detected by an increase in absorbance at 405 nm following the hydrolysis of the substrate *p*NPB. Once again supernatants containing the thermophilic enzymes all exhibited clear activity against the model substrate even at the lowest temperature of 30°C, with the most apparent activity being observed by LCC and PHL7 (Fig. 2C). A low activity against *p*NPB was detected for the mesophilic IsPETase derivative FastPETase, which has been engineered to be significantly more active than the wild-type version of the enzyme (27).

### Solvent-pretreated PET shows high availability for enzymatic hydrolysis

Once the ability of the strains for the secretion of active PET hydrolases had been confirmed, we investigated the use of different types of PET as the reaction substrate. The crystallinity of the polymer is a crucial property defining its degradability and only amorphous PET can be efficiently hydrolysed enzymatically (34,35). We investigated four forms of amorphous PET materials, ranging from films to cryomilled and a sieved cryomilled microplastic fraction (< 100 μm), as well as a solvent pre-treated amorphous macroporous material (specific surface of 45 m^2^ g^-1^) to investigate their susceptibility to enzymatic degradation (Fig. 3A). The different types of PET were added to supernatants of *E. coli* producing a selection of the enzymes in the collection. After incubation for 3 weeks at 30°C, the amount of the PET monomer TA released into the supernatant was determined by measuring the absorbance at 240 nm (Fig. 3B).

**Figure 3.**
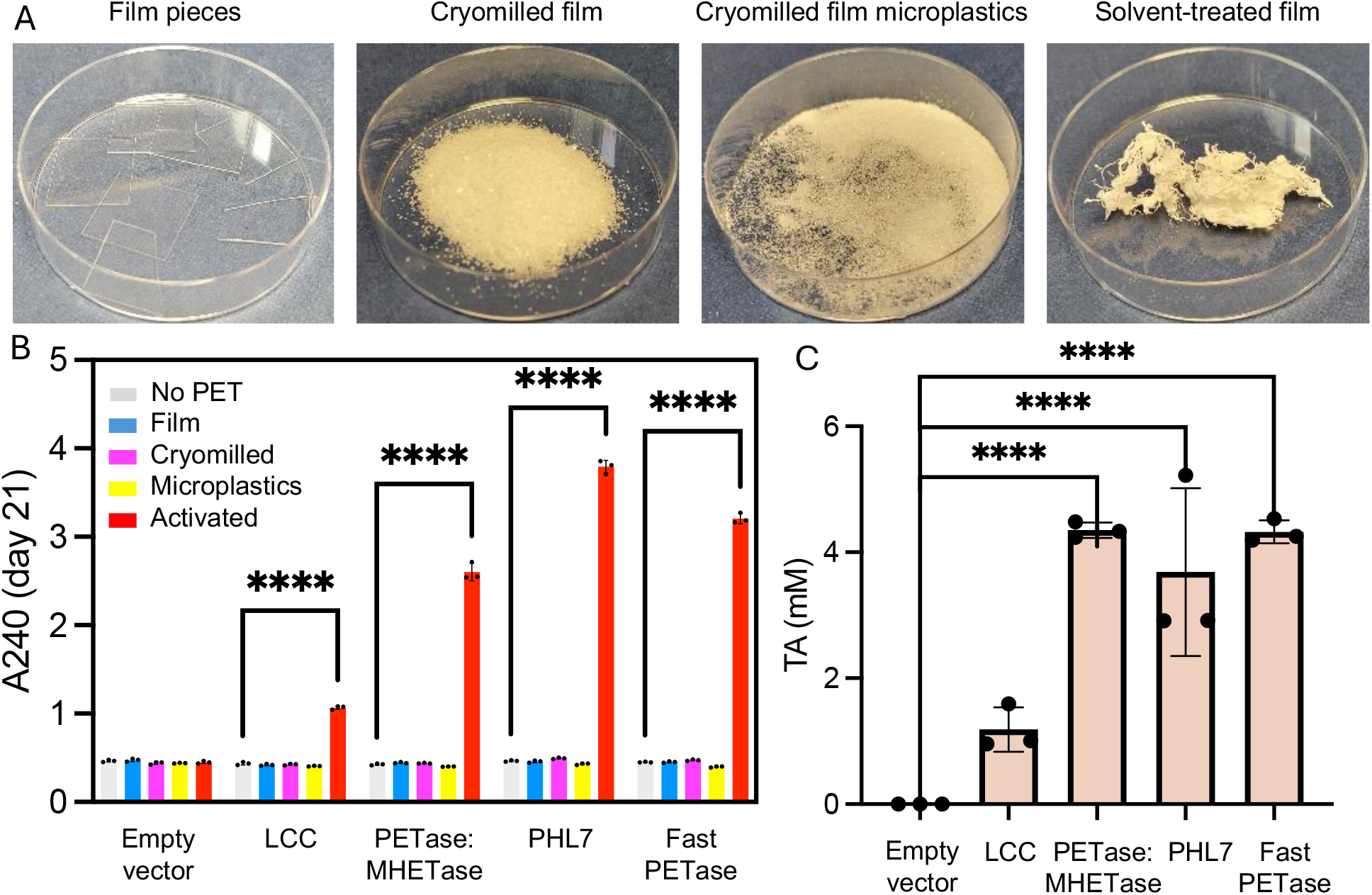
Effect of pretreatments on the availability of PET for enzymatic hydrolysis. (A) Pictures of PET films resulting from the corresponding pre-treatments. (B) TA released from each PET type monitored by measuring the absorbance at 240 nm after incubation at 30°C with cell culture supernatants containing the enzymes shown. (C) Determination of TA by LC-MS in samples of solvent pre-treated PET incubated with the enzymes. Bars and error bars correspond respectively to the mean and SEM of three biological replicates.

Additionally, the presence of TA in supernatants with absorbance above background levels was confirmed by mass spectrometry (MS) (Fig. 3C). Our results show that in the conditions tested, only the solvent-treated PET was hydrolysed enzymatically and, out of the several enzymes tested, PHL7, FastPETase and the chimera MHETase-PETase displayed the highest activity, outperforming LCC. These enzymes generated an amount of TA in the mM range, which in principle should be enough to support microbial growth.

### Utilisation of PET as a microbial feedstock

Our results suggest that the combination of *P. umsongensis* GS as a host for expression, PHL7 as the secreted hydrolase and the use of solvent pre-treated PET would provide the best conditions for PETtrophy. This was evaluated in experiments in which the isolate *P. umsongensis* GS was grown in a minimal medium in the presence of PET and supplemented with different concentrations of LB. This supplementation was required to allow the cells to produce the hydrolytic enzymes as there was no detectable growth in the absence of LB. We determined the contribution of PET hydrolysis to growth by determining the increase in absorbance at 600 nm before and after 1 week of incubation at 30°C of a strain producing PHL7 in the presence or absence of pre-treated PET (Fig. 4A) and the corresponding control with an empty plasmid (Fig. S4). Our results show that the TA alone does not make a difference in the growth of the control strain or can be even detrimental, specially under nutrient-rich conditions (Fig. S4). However, when the strain expresses the PHL7 enzyme that can hydrolyse the polymer, there is a significant difference between the PHL7-producing and the control cultures due to the presence of PET when they are supplemented with the lowest concentration of LB (10%) (p<0.0001), and this difference disappears as the LB concentration is raised, possibly because of the benefits derived from TA as a carbon source diminish when the media becomes richer. PET assimilation was further tested by incubating *P. umsongensis* GS expressing the enzyme PHL7 in the presence or absence of PET for 4 weeks, which resulted in a two-fold increase in the biomass of the culture as a result of the utilisation of the plastic by the strain (Fig. 4B). Taken altogether, these results suggest that PET hydrolysis has a modest but detectable contribution to the growth of *P. umsongensis* GS and confirms the ability of an engineered environmental bacterium to assimilate PET.

**Figure 4.**
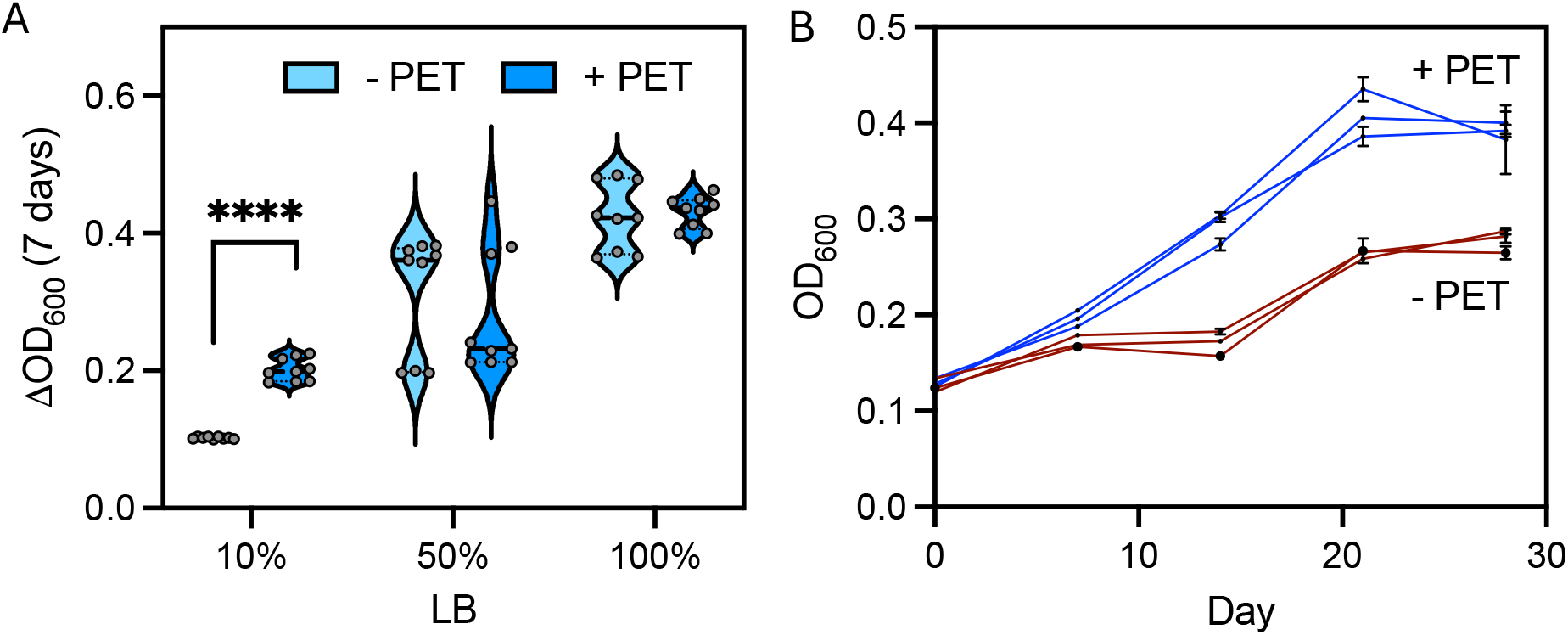
Growth of *P. umsongensis* GS expressing PHL7 with solvent pre-treated PET as a growth substrate. (A) Increment in optical density of the strain expressing PHL7 after 1 week of incubation in the absence (light blue) or presence (dark blue) of solvent pre-treated PET supplemented with different amounts of LB. (B) Growth kinetics of the bacterium expressing PHL7 over 4 weeks in the absence (red) or presence (blue) of activated PET. Results show the individual trajectories of three individual cultures with three replicates each.

## Discussion

In this work we have demonstrated the feasibility of using a whole cell catalyst for the direct assimilation of PET waste plastic under mesophilic conditions. This required the combination of an efficient system for secretion of highly active enzymes against PET expressed in a naturally TA-metabolising bacterium and the use of a PET substrate that had been pre-treated to increase its availability for hydrolysis.

The enzymatic hydrolysis of the polymer was the limiting step in the process. Isolating organisms able to mineralise TA and suitable for genetic manipulation was relatively easy, and we found two strains of *P. umsongensis*, a species that has previously been shown as being able to upcycle TA into the bioplastic polyhydroxyalkanoate (36). However, identifying an enzyme able to support bacterial growth has been a bottleneck in our work and others (20). To overcome this limitation, we tested several of the enzymes described to date, focusing on polyester hydrolases identified both in mesophilic and thermophilic organisms. Thermophilic enzymes have been consistently reported in the literature as endowed with higher activity compared to their mesophilic counterparts many of which are all derived from IsPETase (37). This increased activity at higher temperatures is attributed to the increased chain mobility of the amorphous PET phase as it approaches the polymer’s glass transition (T_g_∼ 70°C), but is, however, incompatible with the optimal growth temperature of microorganisms used in biotechnology that could grow on the resulting monomers. Our results show that thermophilic enzymes present in the supernatants of cell cultures are also more active than IsPETase and its variants at lower temperatures. Out of all the enzymes tested, PHL7 showed the best performance both with PET and the model substrates PCL and *p*NPB in the conditions tested, which include using cell culture medium as the reaction buffer. Interestingly, the activity of PHL7 is known to depend on phosphate concentration that is optimal at 1 M (31). This may explain why it performs well in a relative high-phosphate medium such as M9 containing 0.35 mM phosphate (38). The amount of TA released from PET by this enzyme is in the low mM range and the growth detected by the consolidated environmental isolate is consistent with this concentration and corresponds to half of the maximum biomass obtained when using 10 mM TA as the sole carbon source.

We have shown that the solvent pre-treatment of PET resulted in high hydrolysis rates compared to the same material subjected to other pre-treatments such as cryomilling. This process decreases the crystallinity and increases the porosity of the polymer, which plays an important role in facilitating the access of enzymes to the PET chains.

The modular pipeline of this work successfully demonstrates that active PET hydrolases can be readily expressed in different organisms. However, considering the low PET hydrolysis rates of untreated plastics, whole-cell catalysts are unlikely to efficiently remove these plastics from the environment, particulary microplastics in the sub-millimetre scale. This suggests a need for further enzyme engineering to enhance their activity at the solid-liquid interface of post-consumer PET using methods of screening beyond the conventional colorimetric monitoring of ester hydrolysis using model substrates. It is therefore crucial to develop methods to assess enzymatic efficiency on real-world, heterogeneous materials where the interactions between the enzyme and the polymer interface are more intricate. For example, methods such as impedance spectroscopy (39) can be used for the real-time measurement of plastic film thickness reduction and constitutes a versatile approach applicable to various plastic types.

Although the engineered microorganisms exhibited modest growth on PET, this work lays the groundwork for optimising the process. Engineered PETtrophy could bring additional benefits such as the simplification of the screening of PET hydrolysing enzymes with optimised catalytic properties. Direct selection by growth would facilitate the lab-directed evolution of whole cell organisms, promoting a more efficient PET degradation and aiding the removal of this persistent polymer, while also constituting a significant step towards its upcycling into molecules with added value.

### Experimental Procedures

#### Strain isolation

Soil samples were collected from a range of locations at the University of Surrey campus, UK. Bacterial isolates were cultured from these samples by incubation in 1X M9 minimal medium (5X M9 salts (Sigma) 200 mL/L, 1 M MgSO_4_ 200 µL/L, 1 M CaCl_2_ 100 µL/L, 1,000X trace element solution (ZnSO_4_ · 7H_2_O 0.1 g/L, MnCl_2_ · 4H_2_O 0.03 g/L, H_3_BO_3_ 0.3 g/L, CoCl_2_ · 6H_2_O 0.2 g/L, CuCl_2_ · 2H_2_O 0.01 g/L, NiCl_2_ · 6H_2_O 0.02 g/L, Na_2_MoO_4_ · 2H_2_O 0.03 g/L) 1 mL/L, 10,000X vitamin solution (nicotinic acid 0.01 g/mL, thiamine 0.005 g/L, para-aminobenzoic acid 0.001 g/mL, biotin 0.1 mg/mL) 100 µL/L) supplemented with 10 mM TA. Culturable isolates were streaked to monoculture and maintained on LB agar at 30°C.

#### Whole genome sequencing and analysis

Whole-genome sequencing of environmental isolates was conducted by MicrobesNG, Birmingham, United Kingdom. Briefly, cells were recovered from a LB plate containing pure cultures of each isolate and transferred to a tube containing DNA/RNA Shield (Zymo Research, USA) following MicrobesNG strain submission procedures. 5 to 40 μL of the suspension were lysed with 120 μL of TE buffer containing lysozyme (final concentration 0.1 mg/mL) and RNase A (ITW Reagents, Barcelona, Spain; final concentration 0.1 mg/mL), incubated for 25 min at 37°C. Proteinase K (VWR Chemicals, Ohio, USA) (final concentration 0.1mg/mL) and SDS (Sigma-Aldrich, Missouri, USA) (final concentration 0.5% v/v) were added and incubated for 5 min at 65°C. Genomic DNA was purified using an equal volume of SPRI beads and resuspended in EB buffer (Qiagen, Germany). DNA was quantified with the Quant-iT dsDNA HS kit (ThermoFisher Scientific, United Kingdom) assay in an Eppendorf AF2200 plate reader (Eppendorf UK Ltd, United Kingdom).

Genomic DNA libraries were prepared using the Nextera XT Library Prep Kit (Illumina, San Diego, USA) following the manufacturer’s protocol with the following modifications: input DNA was increased 2-fold, and PCR elongation time was increased to 45 s. DNA quantification and library preparation were carried out on a Hamilton Microlab STAR automated liquid handling system (Hamilton Bonaduz AG, Switzerland). Pooled libraries were quantified using the Kapa Biosystems Library Quantification Kit for Illumina. Libraries were sequenced using Illumina sequencers (HiSeq/NovaSeq) using a 250 bp paired end protocol. Reads were adapter trimmed using Trimmomatic 0.30 with a sliding window quality cutoff of Q15 (40). De novo assembly was performed on samples using SPAdes version 3.7 (41), and contigs were annotated using Prokka 1.11 (42).16S rRNA regions were used for sequence identification using the NCBI BLAST blastn search tool.

Assembled genomes provided by MicrobesNG were annotated using Rapid Annotation using Subsystem Technology (RAST) to predict CDS regions and protein functions. The NCBI BLAST blastp tool was used to align query sequences of TA metabolism genes to predicted protein sequences found in the environmental isolates. The genomes were uploaded to the NCBI’s GenBank with the following accession numbers: JALPTA000000000 (*A. schindleri* LK1), JALPTB000000000 (Rhodococcus sp. HM1), JALPTC000000000 (*P. umsongensis* HM2), JALPTD000000000 (*P. azerbaijanoccidens* SH), JALPTE000000000 (*P. umsongensis* GS), JALPTF000000000 (*R. insidiosa* TS).

#### PET pre-treatment

Cryomilled PET was generated using a SPEX 6875 Freezer/Mill. Amorphous films (Goodfellow, UK) were cut into 1 cm^2^ pieces and subject to two 2 min long cycles of milling at 12 RPS following 4 min of pre-cooling. Cryomilled fragments were sieved through a 150 mesh sieve to recover microplastics smaller than 100 μm.

An amorphous, microporous form of PET was obtained by a solvent pre-treatment. Specifically, 1 g of biaxially oriented PET film (Goodfellow, ES301450, UK) was dissolved in 10 ml of 1,1,1,3,3,3-hexafluoroisopropanol (HFIP) and then added dropwise to 500 mL of isopropanol, pre-cooled to -20°C, under continuous stirring. The resulting precipitate was filtered, washed with isopropanol, and dried at room temperature for 24 h.

#### Cell culturing

Environmental isolates were assayed for growth on the four TA and EG monomers using 1X M9 minimal medium supplemented with 10 mM of each respective monomer. Growth assays were performed at 100 µL volumes in 96-well flat-bottomed microtitre plates using a starting inoculum at OD_600_ ∼ 0.01. Plates were incubated at 30°C in a microplate reader (BMG Clariostar) with continuous orbital shaking and OD_600_ readings were taken at 15 min intervals for 72 h. Routine cell culturing took place in LB medium (Sigma) supplemented with the following antibiotics as required: ampicillin (100 µg/mL), kanamycin (50 µg/mL), chloramphenicol (35 µg/mL), gentamicin (20 µg/mL), streptomycin (100 µg/mL) and tetracycline (10 µg/mL) and 1.5% agar for solid culturing. Supernatants containing secreted enzymes were generated by growing cells for 2 days in LB broth at room temperature. Supernatants were recovered after removing the cells by centrifuging the cultures for 10 min at 12,000 rpm and filtering the supernatants using 0.2 µm cellulose filters (brand Acrodisc). When using PET as growth substrate, 10 mg of activated PET was submerged in 20 mL of M9 medium supplemented with the appropriate concentration of LB. All PET materials were washed with 70% ethanol solution, rinsed with sterilised water and dried prior to their use. Plastic pieces were added to a 5 mL M9 solution supplemented with kanamycin and 0.5 mM benzoate for induction when expressing PET hydrolases. Cultures were incubated at 30°C with orbital shaken at 170 rpm at growth was monitored by taking OD_600_ readings.

#### Plasmid assembly and strain engineering

Genetic constructs for plasmid-based expression of genes of interest were assembled using the broad-host pSEVA238 backbone (43) (Table S3). The plasmid was modified to include a short sequence in the N-terminus of the cloned esterases encoding the peptide leader of the UxpB protein of *P. putida* KT2440. This was achieved by PCR amplifying the secretion leader at the 5’ end of uxpB with primers uxpB-Fwd and uxpB-Rev (Table S4) and conducting Gibson assembly (New England Biolabs, USA) of the resulting fragment into pSEVA238 previously digested with SacI and KpnI. The genes encoding the hydrolases TfCut2, TCur0390, IsPETase, DuraPETase, MHETase and MHETase-PETase (Table S5) were PCR amplified with the appropriate primers (Table S4) using plasmids for intracellular expression as templates (Table S3). The inactive variant TfCut2mut of TfCut2 containing the substitution S187A in the active site was generated by overlapping PCR of the whole plasmid containing the wild type gene using the primers described in Table S4. All esterase coding fragments, including LCC (obtained by digestion of pET20-LCC) and DuraPETase (obtained by DNA synthesis) were cloned in the vector using restriction enzymes BamHI and HindIII followed by ligation with T4 DNA ligase. All Gibson and ligation mixtures were transformed into *E. coli* TOP10 competent cells and selected on LB agar supplemented with kanamycin.

Plasmids were delivered to *E. coli* MG1655 by electroporation in preparation for enzymatic assays. Plasmids were also transferred from *E. coli* to the environmental isolates by triparental mating using *E. coli* DH5α carrying the appropriate plasmids as donors and *E. coli* HB101 pRK600 as a helper (44). Transconjugants were selected in M9 medium containing 5 mM of TA as the only carbon source (selective for the recipient strains) and kanamycin.

#### Esterase enzymatic assays and TA determination

Degradation of PET nanoparticles was conducted in LB-agar plates containing PET nanoparticles prepared as described previously (45). In short, 0.1 g of PET was dissolved in 10 ml HFIP and precipitated by dropping the solution into 100 mL of ice-cold water under stirring and subsequent concentration under vacuum. 100 mg of the nanoparticles were added to 500 mL of molten agar media prior to pouring into Petri dishes. Plates were inoculated with single colonies of esterase producing strains and incubated for 2 days at 30°C followed by 1-day long incubation at 50°C (for assays with thermophilic cutinases). PCL plates were prepared by dissolving 250 mg of PCL in 30 mL of acetone and incubating at 50°C until complete solubilisation. The suspension was mixed slowly with 500 mL of molten LB agar and sonicated in a water bath for 15 minutes before adding any required antibiotics or inducers prior to pouring in petri dishes. Plates were left to dry in a sterile cabinet to allow acetone to evaporate.

Hydrolysis of *p*NPB was conducted using culture supernatants of *E. coli* MG1655 cells (46) grown on LB. Strains containing plasmids for the production and secretion of esterases were cultured to saturation for 14 hours. 1 mL of culture supernatants was collected and cells were removed by centrifugation 10 min at 13,000 rpm followed by filtering with a 0.2 μL syringe filter (Fisher Scientific). Protein concentration in the supernatants was determined using colorimetric Protein Assay (Bio-Rad). All assays were conducted with a volume of freshly prepared supernatant containing 10 ng of total protein (typically in the range of 10 μL) added to a mixture of 80 μL of phosphate buffer 100 mM pH 8.0 and 10 μL of 5 mM *p*NPB in 96-well plates (total volume 100 μL per reaction). Reactions were incubated at 30 or 42°C and monitored measuring the increase in absorbance at 405 nm in a microplate reader (Infinite m200 Pro; Tecan). The absorbance of a control containing *p*NPB in the reaction buffer but lacking culture supernatant was monitored in the same conditions to determine the spontaneous hydrolysis of the substrate. These values were considered background and subtracted from the absorbance of the test samples. The reactions were monitored over time and initial linear absorbance values were fitted by linear regression (GraphPad Prism) in order to obtain the initial reaction velocities.

For enzymatic PET degradation we used either 1 cm^2^ of amorphous films with a thickness of 0.25 mm (Goodfellow, UK), 0.1 g of the same PET cryomilled in advance, 0.1 g of cryomilled PET sieved at <100 µm or 10 mg of solvent pre-treated PET. Plastics were submerged in 1 mL LB culture supernatants containing 5 µg/mL of total protein previously determined in a Bradford Assay (Bio-Rad). Incubations took place at 30°C for 3 weeks. TA released and accumulated in the supernatants was determined spectrophotometrically at 240 nm on a routine basis.

TA was also determined by HPLC-MS. The data were acquired with an Agilent 1290 Infinity II UHPLC coupled to a 6545 LC/Q TOF system. Chromatographic separation was performed with an Agilent InfinityLab Poroshell 120 HILIC-Z (2.1 × 100 mm, 2.7 μm (p/n 675775-924)) column. A concentrated 10x solution consisting of 100 mM ammonium acetate (pH 9.0) in water was prepared to produce mobile phases A and B. Mobile phase A consisted of 10 mM ammonium acetate in water (pH 9) with a 5-μM Agilent InfinityLab Deactivator Additive (p/n 5191-4506), and mobile phase B consisted of 1.0 mM ammonium acetate (pH 9) in 10:90 (v:v) water/acetonitrile with a 5-μM Agilent InfinityLab Deactivator Additive (p/n 5191-4506). The following gradient was applied at a flow rate of 0.5 ml/min: 0 minutes, 100% B; 0-11.5 minutes, 70% B; 11.5-15 minutes, 100% B; 12-15 minutes, 100% B; and 5-minutes of re-equilibration at 100% B. Accurate mass spectrometry was performed using an Agilent Accurate Mass 6545 QTOF apparatus. Dynamic mass axis calibration was achieved by continuous infusion after the chromatography of a reference mass solution using an isocratic pump connected to an ESI ionization source operated in negative-ion mode. The following parameters were used: sheath gas temperature, 300 °C; nebulizer pressure, 40 psig; sheath gas flow, 12 l min-1; capillary voltage, 3000 V; nozzle voltage, 0 V; and fragmentor voltage, 115 V. The data were collected in centroid 4 GHz (extended dynamic range) mode. Data was analysed with the Agilent’s Profinder B.08.00 suite (mass 166.0266).

#### Violacein determination

Violacein present in culture supernatants was determined as described previously (24). Briefly, cells were recovered by centrifugation (13,000 rpm, 10 min) of 1 mL of culture. Violacein was extracted from the cell pellets with absolute ethanol incubating at 95°C for 10 min and it was determined reading absorbance at 570 nm.

#### Statistical analyses

All results show the mean of at least three independent biological replicates with three technical each unless stated otherwise. Evaluation of significant differences was assessed by conducting a one-way ANOVA with a post-hoc Dunnett test or a double tailed t-test when required. Pairwise comparisons were always conducted against the negative control for the experiment. Absence of asterisks in the plots represents no significant differences, while p-values are represented as: **** p<0.0001; *** p<0.001; ** p<0.01; * p<0.1.

## Acknowledgements

The authors are indebted to Dr Tom Bond (University of Surrey, UK) for his support producing cryomilled PET. The authors acknowledge the support received by the Biology and Biotechnology Research Council (BBSRC) through the grant BB/T011289/2 and from the Sächsisches Staatsministerium für Wissenschaft, Kultur und Tourismus (Project No. 100387903) sponsored as part of the ERA-CoBiotech project MIPLACE. The authors would also like to acknowledge the support received from the European Union’s Horizon 2020 research and innovation programme under grant agreement no. 633962 for the project P4SB. U.A. was the recipient of a PhD studentship from the Petroleum Technology Development Fund of Nigeria.

