## Supplementary information for "Degradation of PET plastic with engineered environmental bacteria"

Table S1. Environmental isolates and locus tag of genes forming the *tph* cluster.

| | *tphA1* | *tphA2* | *tphA3* | *tphB* |
|---|---|---|---|---|
| *Rhodococcus* sp. HM1 | M1M07_11370 | M1M07_11355 | M1M07_11360 | M1M07_11365 |
| *Pseudomonas umsongensis* HM2 | M1M10_17875 | M1M10_17895 | M1M10_17890 | M1M10_17885 |
| *Pseudomonas umsongensis* GS | M1M08_06430 | M1M08_06410 | M1M08_06415 | M1M08_06420 |
| *Pseudomonas azerbaijanoccidens* SH | M1M11_02935 | M1M11_02955 | M1M11_02950 | M1M11_02945 |

Table S2. Enzymes used in this study

| **Gene** | **Source** | **Description** | **Reference** |
|---|---|---|---|
| TfCut2-WT | Zimmermann lab | Thermophilic cutinase from *Thermobifida fusca* | (1) |
| TfCut2-mut | *T. fusca* (mutant) | Thermophilic cutinase TfCut2 inactivated by mutation S187A | This work |
| TCur0390 | Zimmermann lab | Thermophilic cutinase from *Thermomonospora curvata* | (2) |
| LCC | Zimmermann lab | Leaf-compost cutinase from metagenomic screening | (3) |
| PHL7 | Zimmermann lab | Thermophilic cutinase from metagenomic screening | (4) |
| IsPETase | *Ideonella sakaiensis* | IsPETase wildtype | (5) |
| IsPETase-modified | Beckham lab via Addgene | IsPETase with mutations W159H-S238F | (6) |
| DuraPETase | DNA synthesis | IsPETase engineered by directed evolution | (7) |
| MHETase | *I. sakaiensis* | IsMHETase wildtype | (5) |
| MHETase-PETase | Beckham lab via Addgene | Translational chimeric fusion of IsMHETase and IsPETase | (6) |
| FAST-PETase | DNA synthesis | IsPETase engineered by AI-assisted directed evolution | (8) |

Table S3. Plasmids used in this study

| Name | Description | Reference |
|---|---|---|
| pSEVA-Hvio | Production of violacein | (9) |
| pET20-TfCut2 | Template for *TfCut2* PCR amplification | (10) |
| pBAD-TOPOT-Cur | Template for *TCur0390* PCR amplification | (2) |
| pET20-LCC | Template for *LCC* cloning | (11) |
| pET20-PHL7 | Template for *PHL7* PCR amplification | (4) |
| pET21b(+)-Is-PETase-W159H-S238F | Template for *PETase-W159H-S238F* PCR amplication | (6) |
| pCJ136 | Template for *MHETase* PCR amplification | (12) |
| pCJ191 | Template for *PETase:MHETase* PCR amplification | (12) |
| pSEVA238 | Cloning backbone, pBBR1 ori, kanamycin resistance, XylS dependent expression | (13) |
| pSEVA238-UxpB | pSEVA238 with *UxpB* secretion leader | This study |
| pSEVA238-TfCut2 | pSEVA238-Uxp expressing Tfcut2 | This study |
| pSEVA238-TfCut2mut | pSEVA238-Uxp expressing Tfcut2mut | This study |
| pSEVA238-TCur | pSEVA238-Uxp expressing TCur | This study |
| pSEVA238-LCC | pSEVA238-Uxp expressing LCC | This study |
| pSEVA238-IsPETase | pSEVA238-Uxp expressing IsPETase | This study |
| pSEVA238-IsPETase_mod | pSEVA238-Uxp expressing IsPETase-W159H-S238F | This study |
| pSEVA238-DuraPETase | pSEVA238-Uxp expressing DuraPETase | This study |
| pSEVA238-MHETase | pSEVA238-Uxp expressing MHETase | This study |
| pSEVA238-MHETase-PETase | pSEVA238-Uxp expressing MHETase-PETase | This study |
| pSEVA238-PHL7 | pSEVA238-Uxp expressing PHL7 | This study |
| pSEVA238-FAST-PETase | pSEVA238-Uxp expressing FAST-PETase | This study |

Table S4. Oligonucleotides used in this study

| Name | Sequence (5'-3') | Use |
|---|---|---|
| F-P<br>R-P | CCTACGGGNGGCWGCAG<br>GACTACHVGGGTATCTAATCC | 16S rRNA sequencing |
| uxpB-F<br><br>uxpB-R | CGGCCGCGCGAATTCGAGCTTTTGTTTAACTTTAAGAAGGAGA<br>TATATCCATGAGTCGAGATACCGGCGGCAACCGAGCGTTC<br>TCTAGAGGATCCCCGGGTACGCAGCCTGTCAGGCCGGT | Cloning of the UxpB secretion leader in pSEVA238 |
| TfCut2-F<br>TfCut2-R | CGCGGATCCGGCAAATCCGTATGAACGTGGTCCGAATCC<br>CGCAAGCTTGTCGACGGAGCTCGAATTCGGATAAAACGGACA | Cloning of *TfCut2* in pSEVA238-UxpB |
| S130A-F<br>S130A-R | ATGACCCATAACTGCCAGAC<br>CTGGCAGTTATGGGTCATGCCATGGGTGGTGGTGGTAGCCTG<br>CG | mutagenesis of TfCut2 |
| TCur-F<br>TCur-R | CGCGGATCCGGCAAATCCGTATCAGC<br>CGCAAGCTTTTACATCGGACAGGTAAAAC | Cloning of *TCur0390* in pSEVA238-UxpB |
| PHL7-F<br>PHL7-R | AAAGGATCCGGCGAACCCGTACGAGCGCGGGCCCGATCCC<br>GGAAAAGCTTTTAGAACGGGCAGGTGGAGCGGTACTCGG | Cloning of *PHL7* in pSEVA238-UxpB |
| Is-F<br>Is-R | TATTATAATTAGGATCCGGCCAACCCCTACGC<br>GCAAGCTTTCAGCTGCAGTTCGC | Cloning of *IsPETase* in pSEVA238-UxpB |
| Ism-F<br>Ism-R | TATTATAATTAGGATCCGAATCCGTATGCGC<br>CGCAAGCTTTCAGGAACAGTTCGC | Cloning of *IsPETase W159H-S238F* in pSEVA238-UxpB |
| FAST-F<br>FAST-R | CCCGGGATCCGGAACCCCTACGCCC<br>AATTAAGCTTTCAGCTGCAGTTCGCGG | Cloning of *FAST-PETase* in pSEVA238-UxpB |
| MHET-F<br>MHET-R | TATTATAATTAGGATCCGTGCGCCGGAGGAG<br>TAATAAGCTTTCACGGTGGAGCGGC | Cloning of *MHETase* in pSEVA238-UxpB |
| MP-F<br>MP-R | TATTATAATTAGGATCCGTGCGCCGGAGGAG<br>CGCAAGCTTTCAGGAACAGTTCGC | Cloning of *MHETase-PETase* in pSEVA238-UxpB |

Table S5. DNA and amino acid sequences of the enzymes used in this work (all native secretion signals were replaced by the secretion leader of the protein UxpB from *P. putida* KT2440 (shown in red). Blue segments denote linker regions.

| | |
|---|---|
| **TfCut2** | |
| nucleotide | ATGAGTCGAGATACCGGCGACAACCTGGACCGGAACCAAAGCGGCAACCTGCCGATGGCCAACGTCATGGACGCCTAC<br>CTGAGCCGCCGCAGCGTGATGCGCGGCAGCCTGGGCGCCGCCATCGCCATGATTGCCGGTACCGGCCTGACAGGCTGC<br>GTACCCGGGGATCCGGCAAATCCGTATGAACGTGGTCCGAATCCGACCGATGCACTGCTGGAAGCACGTAGCGGTCCG<br>TTTAGCGTTAGCGAAGAAAATGTTAGCCGTCTGAGCGCAAGCGGTTTTGGTGGTGGCACCATTTATTATCCGCGTGAA<br>AATAATACCTATGGTGCAGTTGCAATTAGTCCGGGTTATACCGGCACCGAAGCAAGCATTGCATGGCTGGGTGAACGT<br>ATTGCAAGCCATGGTTTTGTTGTGATTACCATTGATACCATTACCACCCTGGATCAGCCGGATAGCCGTGCAGAACAG<br>CTGAATGCAGCACTGAATCATATGATTAATCGTGCAAGCAGCACCGTTCGTAGCCGTATTGATAGCAGCCGTCTGGCA<br>GTTATGGGTCATAGCATGGGTGGTGGTGGTAGCCTGCGTCTGGCAAGCCAGCGTCCGGATCTGAAAGCAGCAATTCCG<br>CTGACCCCGTGGCATCTGAATAAAAATTGGAGCAGCGTTACCGTTCCGACCCTGATTATTGGTGCAGATCTGGATACC<br>ATTGCACCGGTTGCAACCCATGCAAAACCGTTTTATAATAGCCTGCCGAGCAGCATTAGCAAAGCATATCTGGAACTG<br>GATGGTGCAACCCATTTTGCACCGAATATTCCGAATAAAATTATTGGCAAATATAGCGTGGCCTGGCTGAAACGTTTT<br>GTTGATAATGATACCCGCTATACCCAGTTTCTGTGTCCGGGTCCGCGTGATGGTCTGTTTGGTGAAGTTGAAGAATAT<br>CGTAGCACGTGTCCGTTTTATCCGAATTCGAGCTCCGTCGACAAGCTTGCGGCCGCGTCGTGA |
| protein | MSRDTGDNLDRNQSGNLPMANVMDAYLSRRSVMRGSLGAAIAMIAGTGLTGCVPGDPANPYERGPNPTDALLEARSGP<br>FSVSEENVSRLSASGFGGGTIYYPRENNTYGAVAISPGYTGTEASIAWLGERIASHGFVVITIDTITTLDQPDSRAEQ<br>LNAALNHMINRASSTVRSRIDSSRLAVMGHSMGGGGSLRLASQRPDLKAAIPLTPWHLNKNWSSVTVPTLIIGADLDT<br>IAPVATHAKPFYNSLPSSISKAYLELDGATHFAPNIPNKIIGKYSVAWLKRFVDNDTRYTQFLCPGPRDGLFGEVEEY<br>RSTCPFYPNSSSVDKLAAAS |
| **TCur0390** | |
| nucleotide | ATGAGTCGAGATACCGGCGACAACCTGGACCGGAACCAAAGCGGCAACCTGCCGATGGCCAACGTCATGGACGCCTAC<br>CTGAGCCGCCGCAGCGTGATGCGCGGCAGCCTGGGCGCCGCCATCGCCATGATTGCCGGTACCGGCCTGACAGGCTGC<br>GTACCCGGGGATCCGGCAAATCCGTATCAGCGTGGTCCGAATCCGACCGAAGCAAGCATTACCGCAGCACGTGGTCCG<br>TTTAATACCGCAGAAATTACCGTTAGCCGTCTGAGCGTTTCAGGTTTTGGTGGCAAAATTTATTATCCGACCACC<br>ACCTCTGAAGGCACCTTTGGTGCAATTGCAATTAGTCCGGGTTTTACCGCATATTGGAGCAGCCTGGAATGGCTGGGT<br>CATCGTCTGGCAAGCCAGGGTTTTGTTGTTATTGGTATTGAAACCAATACCACCCTGGATCAGCCGGATCAGCGTGGA<br>CAGCAACTGCTGGCAGCACTGGATTATCTGACCCAGCGTAGCGCAGTTCGTGATCGTGTTGATGCAAGCCGTCTGGCA<br>GTTGCAGGTCATAGCATGGGTGGTGGTGGTAGCCTGGAAGCAGCAAAAGCACGTACCAGCCTGAAAGCAGCCATTCCG<br>CTGGCACCGTGGAATCTGGATAAAACCTGGCCTGAAGTTCGTACTCCGACCCTGATTATTGGTGGTGAACTGGATGCA<br>GTTGCACCGGTTGCTACCCATAGCATTCCGTTTTATAATAGCCTGAGCAATGCACCGGAAAAAGCATATCTGGAACTG<br>GATAATGCCAGCCATTTTTTTCCGAATATTACCAATACCCAGATGGCCAAATATATGATTGCCTGGATGAAACGCTTT<br>ATTGATGATGATACCCGCTATACCCAGTTTCTGTGTCCGCCTCCGAGCACCGGTCTGCTGAGCGATTTTAGTGATGCA<br>CGTTTTACCTGTCCGATGTAA |
| protein | MSRDTGDNLDRNQSGNLPMANVMDAYLSRRSVMRGSLGAAIAMIAGTGLTGCVPGDPANPYQRGPNPTEASITAARGP<br>FNTAEITVSRLSVSGFGGGKIYYPTTTSEGTFGAIAISPGFTAYWSSLEWLGHRLASQGFVVIGIETNTTLDQPDQRG<br>QQLLAALDYLTQRSAVRDRVDASRLAVAGHSMGGGGSLEAAKARTSLKAAIPLAPWNLDKTWPEVRTPTLIIGGELDA<br>VAPVATHSIPFYNSLSNAPEKAYLELDNASHFFPNITNTQMAKYMIAWMKRFIDDDTRYTQFLCPPPSTGLLSDFSDA<br>RFTCPM |
| **LCC** | |
| nucleotide | ATGAGTCGAGATACCGGCGACAACCTGGACCGGAACCAAAGCGGCAACCTGCCGATGGCCAACGTCATGGACGCCTAC<br>CTGAGCCGCCGCAGCGTGATGCGCGGCAGCCTGGGCGCCGCCATCGCCATGATTGCCGGTACCGGCCTGACAGGCTGC<br>GTACCCGGGGATCCGTCTAACCCGTACCAGCGTGGTCCGAACCCGACCCGTTCTGCTCTGACCGCTGACGGTCCGTTC<br>TCTGTTGCTACCTACACCGTTTCTCGTCTGTCTGTTTCTGGTTTCGGTGGTGGTGTTATCTACTACCCGACCGGTACC<br>TCTCTGACCTTCGGTGGTATCGCTATGTCTCCGGGTTACACCGCTGACGCTTCTTCTCTGGCTTGGCTGGGTCGTCGT<br>CTGGCTTCTCACGGTTTCGTTGTTCTGGTTATCAACACCAACTCTCGTTTCGACTACCCGGACTCTCGTGCTTCTCAG<br>CTGTCTGCTGCTCTGAACTACCTGCGTACCTCTTCTCCGTCTGCTGTTCGTGCTCGTCTGGACGCTAACCGTCTGGCT<br>GTTGCTGGTCACTCTATGGGTGGTGGTGGTACCCTGCGTATCGCTGAACAGAACCCGTCTCTGAAAGCTGCTGTTCCG<br>CTGACCCCGTGGCACACCGACAAAACCTTCAACACCTCTGTTCCGGTTCTGATCGTTGGTGCTGAAGCTGACACCGTT<br>GCTCCGGTTTCTCAGCACGCTATCCCGTTCTACCAGAACCTGCCGTCTACCACCCCGAAAGTTTACGTTGAACTGGAC<br>AACGCTTCTCACTTCGCTCCGAACTCTAACAACGCTGCTATCTCTGTTTACACCATCTCTTGGATGAAACTGTGGGTT<br>GACAACGACACCCGTTACCGTCAGTTCCTGTGCAACGTTAACGACCCGGCTCTGTCTGACTTCCGTACCAACAACCGT<br>CACTGCCAGTAA |
| protein | MSRDTGDNLDRNQSGNLPMANVMDAYLSRRSVMRGSLGAAIAMIAGTGLTGCVPGDPSNPYQRGPNPTRSALTADGPF<br>SVATYTVSRLSVSGFGGGVIYYPTGTSLTFGGIAMSPGYTADASSLAWLGRRLASHGFVVLVINTNSRFDYPDSRASQ<br>LSAALNYLRTSSPSAVRARLDANRLAVAGHSMGGGGTLRIAEQNPSLKAAVPLTPWHTDKTFNTSVPVLIVGAEADTV<br>APVSQHAIPFYQNLPSTTPKVYVELDNASHFAPNSNNAAISVYTISWMKLWVDNDTRYRQFLCNVNDPALSDFRTNNR<br>HCQ |
| **PHL7** | |
| nucleotide | ATGAGTCGAGATACCGGCGACAACCTGGACCGGAACCAAAGCGGCAACCTGCCGATGGCCAACGTCATGGACGCCTAC<br>CTGAGCCGCCGCAGCGTGATGCGCGGCAGCCTGGGCGCCGCCATCGCCATGATTGCCGGTACCGGCCTGACAGGCTGC<br>GTACCCGGGGATCCGGCGAACCCGTACGAGCGCGGGCCCGATCCCACCGAGTCGAGCATCGAGGCCGTCCGCGGGCCG<br>TTCGCCGTGGCCCAGACGACGGTGTCGAGGCTCCAGGCCGACGGCTTCGGCGGCGGACCATCTACTACCCGACCGAC<br>ACGAGCCAGGGCACCTTCGGTGCGGTGGCGATCTCGCCGGGGTTCACGGCGGGCCAGGAGAGCATCGCCTGGCTCGGC<br>CCCCGCATCGCGTCGCAGGGCTTCGTGGTGATCACGATCGACACGATCACGCGCCTCGACCAGCCCGACAGCCGGGGT<br>CGCCAGCTGCAGGCCGCGCTCGACCACCTGCGCACCAACAGCGTCGTGCGCAACCGGATCGACCCGAACCGGATGGCG<br>GTCATGGGCCACTCGATGGGCGGCGGCGGGGCGCTGTCCGCCGCGGCGAACAACACGAGCCTCGAGGCCGCCATCCCG<br>CTGCAGGGCTGGCACACCCGGAAGAACTGGTCGAGCGTGCGGACGCCGACCCTGGTGGTCGGGGCCCAGCTCGACACC<br>ATCGCGCCGGTGAGCTCGCACTCGGAGGCCTTCTACAACAGCCTGCCGAGCGACCTCGACAAGGCGTACATGGAGCTC<br>CGCGGGGCCAGCCACCTCGTGTCGAACACGCCCGACACGACGACCGCGAAGTACAGCATCGCCTGGCTCAAGCGGTTC |

| | |
|---|---|
| | GTCGACGACGACCTCCGCTACGAGCAGTTCCTGTGCCCGGCGCCGGACGACTTCGCGATCTCCGAGTACCGCTCCACC<br>TGCCCGTTCTAA |
| protein | MSRDTGDNLDRNQSGNLPMANVMDAYLSRRSVMRGSLGAAIAMIAGTGLTGCVPGDPANPYERGPDPTESSIEAVRGP<br>FAVAQTTVSRLQADGFGGGTIYYPTDTSQGTFGAVAISPGFTAGQESIAWLGPRIASQGFVVITIDTITRLDQPDSRG<br>RQLQAALDHLRTNSVVRNRIDPNRMAVMGHSMGGGGALSAAANNTSLEAAIPLQGWHTRKNWSSVRTPTLVVGAQLDT<br>IAPVSSHSEAFYNSLPSDLDKAYMELRGASHLVSNTPDTTTAKYSIAWLKRFVDDDLRYEQFLCPAPDDFAISEYRST<br>CPF |
| **IsPETase** | |
| nucleotide | ATGAGTCGAGATACCGGCGACAACCTGGACCGGAACCAAAGCGGCAACCTGCCGATGGCCAACGTCATGGACGCCTAC<br>CTGAGCCGCCGCAGCGTGATGCGCGGCAGCCTGGGCGCCGCCATCGCCATGATTGCCGGTACCGGCCTGACAGGCTGC<br>GTACCCGGGGATCCGGCCAACCCCTACGCCCGCGGCCCGAACCCGACAGCCGCCTCACTCGAAGCCAGCGCCGGCCCG<br>TTCACCGTGCGCTCGTTCACCGTGAGCCGCCCGAGCGGCTACGGCGCCGGCACCGTGTACTACCCCACCAACGCCGGC<br>GGCACCGTGGGCGCCATCGCCATCGTGCCGGGCTACACCGCGCGCCAGTCGAGCATCAAATGGTGGGGCCCGCGCCTG<br>GCCTCGCACGGCTTCGTGGTCATCACCATCGACACCAACTCCACGCTCGACCAGCCGTCCAGCCGCTCGTCGCAGCAG<br>ATGGCCGCGCTGCGCCAGGTGGCCTCGCTCAACGGCACCAGCAGCAGCCCGATCTACGGCAAGGTCGACACCGCCCGC<br>ATGGGCGTGATGGGCTGGTCGATGGGCGGTGGCGGCTCGCTGATCTCGGCGGCCAACAACCCGTCGCTGAAAGCCGCG<br>GCGCCGCAGGCCCCGTGGGACAGCTCGACCAACTTCTCGTCGGTCACCGTGCCCACGCTGATCTTCGCCTGCGAGAAC<br>GACAGCATCGCCCCGGTCAACTCGTCCGCCCTGCCGATCTACGACAGCATGTCGCGCAATGCGAAGCAGTTCCTCGAG<br>ATCAACGGTGGCTCGCACTCCTGCGCCAACAGCGGCAACAGCAACCAGGCGCTGATCGGCAAGAAGGGCGTGGCCTGG<br>ATGAAGCGCTTCATGGACAACGACACGCGCTACTCCACCTTCGCCTGCGAGAACCCGAACAGCACCCGCGTGTCGGAC<br>TTCCGCACCGCGAACTGCAGCTGA |
| protein | MSRDTGDNLDRNQSGNLPMANVMDAYLSRRSVMRGSLGAAIAMIAGTGLTGCVPGDPANPYARGPNPTAASLEASAGP<br>FTVRSFTVSRPSGYGAGTVYYPTNAGGTVGAIAIVPGYTARQSSIKWWGPRLASHGFVVITIDTNSTLDQPSSRSSQQ<br>MAALRQVASLNGTSSSPIYGKVDTARMGVMGWSMGGGGSLISAANNPSLKAAAPQAPWDSSTNFSSVTVPTLIFACEN<br>DSIAPVNSSALPIYDSMSRNAKQFLEINGGSHSCANSGNSNQALIGKKGVAWMKRFMDNDTRYSTFACENPNSTRVSD<br>FRTANCS |
| **IsPETase-W159H-S238F** | |
| nucleotide | ATGAGTCGAGATACCGGCGACAACCTGGACCGGAACCAAAGCGGCAACCTGCCGATGGCCAACGTCATGGACGCCTAC<br>CTGAGCCGCCGCAGCGTGATGCGCGGCAGCCTGGGCGCCGCCATCGCCATGATTGCCGGTACCGGCCTGACAGGCTGC<br>GTACCCGGGGATCCGAATCCGTATGCGCGCGGCCCCAACCCTACCGCCGCCTCGTTGGAAGCCAGCGCGGGACCCTTT<br>ACCGTTCGTAGCTTTACCGTTAGCCGTCCGTCCGGATATGGTGCAGGGACCGTCTATTACCCAACCAATGCAGGCGGC<br>ACCGTTGGCGCGATTGCAATCGTCCCCGGGTACACCGCGCGTCAAAGCAGCATTAAGTGGTGGGGTCCGCGCTTAGCT<br>AGCCATGGCTTTGTGGTTATTACCATCGATACGAACAGCACTCTAGACCAGCCCAGCAGCCGTAGCTCGCAACAGATG<br>GCCGCGCTTCGTCAAGTTGCGAGCTTGAACGGGACCAGCAGTAGCCCGATTTACGGAAAGGTCGATACTGCCCGCATG<br>GGTGTGATGGGCCACTCAATGGGGGGCGGCGGTTCACTTATTAGCGCCGCGAACAACCCGAGTTTAAAAGCAGCGGCA<br>CCGCAGGCGCCATGGGACTCTTCAACCAACTTCAGCAGTGTTACCGTGCCGACGCTGATTTTCGCGTGCGAGAATGAT<br>AGCATTGCACCGGTGAACAGCAGCGCGCTGCCGATTTATGATAGCATGTCCCGCAACGCAAAACAGTTTCTGGAAATT<br>AACGGCGGTAGCCACTTCTGTGCCAACTCTGGGAACAGCAACCAGGCACTGATCGGAAAAAAAGGGGTTGCATGGATG<br>AAACGATTCATGGATAATGACACCCGTTACTCAACCTTCGCCTGTGAGAATCCCAACAGCACACGCGTGTCGGATTTT<br>CGCACCGCGAACTGTTCCTGA |
| amino acid | MSRDTGDNLDRNQSGNLPMANVMDAYLSRRSVMRGSLGAAIAMIAGTGLTGCVPGDPNPYARGPNPTAASLEASAGPF<br>TVRSFTVSRPSGYGAGTVYYPTNAGGTVGAIAIVPGYTARQSSIKWWGPRLASHGFVVITIDTNSTLDQPSSRSSQQM<br>AALRQVASLNGTSSSPIYGKVDTARMGVMGHSMGGGGSLISAANNPSLKAAAPQAPWDSSTNFSSVTVPTLIFACEND<br>SIAPVNSSALPIYDSMSRNAKQFLEINGGSHFCANSGNSNQALIGKKGVAWMKRFMDNDTRYSTFACENPNSTRVSDF<br>RTANCS |
| **DuraPETase** | |
| nucleotide | ATGAGTCGAGATACCGGCGACAACCTGGACCGGAACCAAAGCGGCAACCTGCCGATGGCCAACGTCATGGACGCCTAC<br>CTGAGCCGCCGCAGCGTGATGCGCGGCAGCCTGGGCGCCGCCATCGCCATGATTGCCGGTACCGGCCTGACAGGCTGC<br>GTACCCGGGCCGGCGAACCCGTATGCGCGCGGCCCGAACCCGACCGCGGCGAGCCTGGAAGCGAGCGCGGGCCCGTTT<br>ACCGTGCGCAGCTTTACCGTGAGCCGCCCGAGCGGCTATGGCGCGGGCACCGTGTATTATCCGACCAACGCGGGCGGC<br>ACCGTGGGCGCGATTGCGATTGTGCCGGGCTATACCGCGCGCCAGAGCAGCATTAAATGGTGGGGCCCGCGCCTGGCG<br>AGCCATGGCTTTGTGGTGATTACCATTGATACCAACAGCACCTTTGATTATCCGAGCAGCCGCAGCAGCCAGCAGATG<br>GCGGCGCTGCGCCAGGTGGCGAGCCTGAACGGCGATAGCAGCAGCCCGATTTATGGCAAAGTGGATACCGCGCGCATG<br>GGCGTGATGGGCCATAGCATGGGCGGCGGCGCGAGCCTGCGCAGCGCGGCGAACAACCCGAGCCTGAAAGCGGCGATT<br>CCGCAGGCGCCGTGGGATAGCAGCACCAACTTTAGCAGCGTGACCGTGCCGACCCTGATTTTTGCGTGCGAAAACGAT<br>AGCATTGCGCCGGTGAACAGCCATGCGCTGCCGATTTATGATAGCATGAGCCGCAACGCGAAACAGTTTCTGGAAATT<br>AACGGCGGCAGCCATAGCTGCGCGAACAGCGGCAACAGCAACCAGGCGCTGATTGGCAAAAAAGGCGTGGCGTGGATG<br>AAACGCTTTATGGATAACGATACCCGCTATAGCACCTTTGCGTGCGAAAACCCGAACAGCACCGCGGTGAGCGATTTT<br>CGCACCGCGAACTGCAGCTGA |
| amino acid | MSRDTGDNLDRNQSGNLPMANVMDAYLSRRSVMRGSLGAAIAMIAGTGLTGCVPGPANPYARGPNPTAASLEASAGPF<br>TVRSFTVSRPSGYGAGTVYYPTNAGGTVGAIAIVPGYTARQSSIKWWGPRLASHGFVVITIDTNSTFDYPSSRSSQQM<br>AALRQVASLNGDSSSPIYGKVDTARMGVMGHSMGGGASLRSAANNPSLKAAIPQAPWDSSTNFSSVTVPTLIFACEND<br>SIAPVNSHALPIYDSMSRNAKQFLEINGGSHSCANSGNSNQALIGKKGVAWMKRFMDNDTRYSTFACENPNSTAVSDF<br>RTANCS |
| **FastPETase** | |
| nucleotide | ATGAGTCGAGATACCGGCGACAACCTGGACCGGAACCAAAGCGGCAACCTGCCGATGGCCAACGTCATGGACGCCTAC<br>CTGAGCCGCCGCAGCGTGATGCGCGGCAGCCTGGGCGCCGCCATCGCCATGATTGCCGGTACCGGCCTGACAGGCTGC<br>GTACCCGGGGATCCGAACCCCTACGCCCGCGGCCCGAACCCGACAGCCGCCTCACTCGAAGCCAGCGCCGGCCCGTTC<br>ACCGTGCGCTCGTTCACCGTGAGCCGCCCGAGCGGCTACGGCGCCGGCACCGTGTACTACCCCACCAACGCCGGCGGC<br>ACCGTGGGCGCCATCGCCATCGTGCCGGGCTACACCGCGCGCCAGTCGAGCATCAAATGGTGGGGCCCGCGCCTGGCC<br>TCGCACGGCTTCGTGGTCATCACCATCGACACCAACTCCACGCTCGACCAGCCGGAAAGCCGCTCGTCGCAGCAGATG<br>GCCGCGCTGCGCCAGGTGGCCTCGCTCAACGGCACCAGCAGCAGCCCGATCTACGGCAAGGTCGACACCGCCCGCATG<br>GGCGTGATGGGCTGGTCGATGGGCGGTGGCGGCTCGCTGATCTCGGCGGCCAACAACCCGTCGCTGAAAGCCGCGGCG<br>CCGCAGGCCCCGTGGCATAGCTCGACCAACTTCTCGTCGGTCACCGTGCCCACGCTGATCTTCGCCTGCGAGAACGAC<br>AGCATCGCCCCGGTCAACTCGTCCGCCCTGCCGATCTACGACAGCATGTCGCAGAATGCGAAGCAGTTCCTCGAGATC |

| | |
|---|---|
| | AAAGGTGGCTCGCACTCCTGCGCCAACAGCGGCAACAGCAACCAGGCGCTGATCGGCAAGAAGGGCGTGGCCTGGATG<br>AAGCGCTTCATGGACAACGACACGCGCTACTCCACCTTCGCCTGCGAGAACCCGAACAGCACCGCGGTGTCGGACTTC<br>CGCACCGCGAACTGCAGCTGA |
| amino acid | MSRDTGDNLDRNQSGNLPMANVMDAYLSRRSVMRGSLGAAIAMIAGTGLTGCVPGDPNPYARGPNPTAASLEASAGPF<br>TVRSFTVSRPSGYGAGTVYYPTNAGGTVGAIAIVPGYTARQSSIKWWGPRLASHGFVVITIDTNSTLDQPESRSSQQM<br>AALRQVASLNGTSSSPIYGKVDTARMGVMGWSMGGGGSLISAANNPSLKAAAPQAPWHSSTNFSSVTVPTLIFACEND<br>SIAPVNSSALPIYDSMSQNAKQFLEIKGGSHSCANSGNSNQALIGKKGVAWMKRFMDNDTRYSTFACENPNSTAVSDF<br>RTANCS |
| **MHETase** | |
| nucleotide | ATGAGTCGAGATACCGGCGACAACCTGGACCGGAACCAAAGCGGCAACCTGCCGATGGCCAACGTCATGGACGCCTAC<br>CTGAGCCGCCGCAGCGTGATGCGCGGCAGCCTGGGCGCCGCCATCGCCATGATTGCCGGTACCGGCCTGACAGGCTGC<br>GTACCCGGGGATCCGTGCGCCGGAGGAGGTTCCACTCCTCTGCCTCTACCGCAGCAGCAGCCGCCTCAGCAGGAACCG<br>CCACCTCCTCCTGTTCCGCTAGCCAGTCGCGCCGCGTGTGAGGCGCTCAAAGATGGTAATGGCGACATGGTTTGGCCG<br>AATGCCGCCACGGTTGTAGAGGTTGCAGCCTGGCGTGATGCAGCACCGGCCACGGCATCAGCCGCAGCCCTGCCGGAG<br>CATTGCGAAGTATCAGGCGCGATTGCCAAGCGTACTGGGATTGATGGGTACCCGTATGAAATTAAGTTTCGCCTGCGC<br>ATGCCCGCTGAGTGGAACGGCCGTTTTTTCATGGAGGGTGGCAGTGGTACGAACGGCTCTCTCTCAGCGGCGACCGGA<br>AGTATCGGCGGCGGTCAGATCGCCTCAGCGCTGAGTCGTAACTTTGCAACAATTGCTACCGACGGAGGACATGACAAT<br>GCGGTGAATGATAATCCGGATGCGCTCGGTACCGTCGCATTTGGTCTCGATCCCCAGGCACGCTTAGACATGGGCTAC<br>AACTCCTATGATCAGGTGACTCAGGCCGGCAAAGCCGCCGTTGCACGCTTTTATGGTCGCGCAGCCGACAAGAGCTAC<br>TTCATCGGCTGTTCGGAGGGCGGCCGCGAGGGCATGATGCTGTCCCAGCGCTTTCCATCACATTACGATGGCATTGTG<br>GCGGGCGCACCGGGATATCAGTTGCCGAAGGCCGGAATTAGTGGCGCGTGGACCACCCAGAGCTTAGCGCCCGCCGCC<br>GTTGGCCTGGATGCCCAGGGAGTGCCGCTGATTAATAAGAGCTTTTCTGACGCAGACCTCCATTTACTGTCGCAGGCG<br>ATTCTCGGAACATGCGACGCCTTGGATGGCCTGGCCGACGGCATCGTTGACAACTACCGAGCGTGCCAAGCGGCTTTT<br>GATCCGGCGACTGCAGCCAACCCAGCGAATGGCCAAGCCCTGCAGTGCGTGGGCGCAAAGACAGCCGATTGCTTATCG<br>CCCGTCCAAGTTACGGCGATTAAACGAGCGATGGCCGGTCCGGTAAATAGCGCGGGTACGCCGTTATATAATAGATGG<br>GCCTGGGACGCAGGTATGAGCGGTCTTAGTGGTACCACTTACAATCAGGGTTGGCGCAGCTGGTGGCTGGGATCGTTT<br>AACAGCTCGGCGAATAACGCACAACGTGTATCTGGTTTCTCAGCGCGGAGCTGGCTGGTGGACTTTGCTACCCCGCCG<br>GAGCCGATGCCCATGACCCAAGTCGCCGCCCGTATGATGAAATTTGATTTCGATATCGATCCTCTGAAAATATGGGCT<br>ACTTCGGGCCAATTTACCCAGAGTAGTATGGACTGGCACGGTGCCACTAGCACCGACCTTGCTGCCTTTCGGGACCGC<br>GGCGGTAAAATGATTCTGTATCACGGAATGAGCGATGCCGCATTCTCTGCACTAGATACAGCAGATTATTATGAACGC<br>CTGGGTGCCGCAATGCCGGGCGCCGCGGGCTTTGCTCGTCTGTTCTTGGTTCCGGGAATGAACCATTGCTCCGGGGGT<br>CCAGGTACCGACCGCTTTGATATGCTAACACCGTTAGTTGCATGGGTTGAACGTGGGGAAGCCCCTGACCAAATTAGC<br>GCCTGGAGCGGCACCCCCGGCTACTTTGGTGTGGCCGCCCCGCACTCGACCGTTATGTCCCTATCCGCAGATTGCGCGC<br>TATAAGGGATCAGGCGATATCAATACCGAAGCAAATTTTGCGTGTGCCGCTCCACCGTGA |
| amino acid | MSRDTGDNLDRNQSGNLPMANVMDAYLSRRSVMRGSLGAAIAMIAGTGLTGCVPGDPCAGGGSTPLPLPQQQPPQQEP<br>PPPPVPLASRAACEALKDGNGDMVWPNAATVVEVAAWRDAAPATASAAALPEHCEVSGAIAKRTGIDGYPYEIKFRLR<br>MPAEWNGRFFMEGGSGTNGSLSAATGSIGGGQIASALSRNFATIATDGGHDNAVNDNPDALGTVAFGLDPQARLDMGY<br>NSYDQVTQAGKAAVARFYGRAADKSYFIGCSEGGREGMMLSQRFPSHYDGIVAGAPGYQLPKAGISGAWTTQSLAPAA<br>VGLDAQGVPLINKSFSDADLHLLSQAILGTCDALDGLADGIVDNYRACQAAFDPATAANPANGQALQCVGAKTADCLS<br>PVQVTAIKRAMAGPVNSAGTPLYNRWAWDAGMSGLSGTTYNQGWRSWWLGSFNSSANNAQRVSGFSARSWLVDFATPP<br>EPMPMTQVAARMMKFDFDIDPLKIWATSGQFTQSSMDWHGATSTDLAAFRDRGGKMILYHGMSDAAFSALDTADYYER<br>LGAAMPGAAGFARLFLVPGMNHCSGGPGTDRFDMLTPLVAWVERGEAPDQISAWSGTPGYFGVAARTRPLCPYPQIAR<br>YKGSGDINTEANFACAAPP |
| **MHETase-PETase** | |
| nucleotide | ATGAGTCGAGATACCGGCGACAACCTGGACCGGAACCAAAGCGGCAACCTGCCGATGGCCAACGTCATGGACGCCTAC<br>CTGAGCCGCCGCAGCGTGATGCGCGGCAGCCTGGGCGCCGCCATCGCCATGATTGCCGGTACCGGCCTGACAGGCTGC<br>GTACCCGGGGATCCGTGCGCCGGAGGAGGTTCCACTCCTCTGCCTCTACCGCAGCAGCAGCCGCCTCAGCAGGAACCG<br>CCACCTCCTCCTGTTCCGCTAGCCAGTCGCGCCGCGTGTGAGGCGCTCAAAGATGGTAATGGCGACATGGTTTGGCCG<br>AATGCCGCCACGGTTGTAGAGGTTGCAGCCTGGCGTGATGCAGCACCGGCCACGGCATCAGCCGCAGCCCTGCCGGAG<br>CATTGCGAAGTATCAGGCGCGATTGCCAAGCGTACTGGGATTGATGGGTACCCGTATGAAATTAAGTTTCGCCTGCGC<br>ATGCCCGCTGAGTGGAACGGCCGTTTTTTCATGGAGGGTGGCAGTGGTACGAACGGCTCTCTCTCAGCGGCGACCGGA<br>AGTATCGGCGGCGGTCAGATCGCCTCAGCGCTGAGTCGTAACTTTGCAACAATTGCTACCGACGGAGGACATGACAAT<br>GCGGTGAATGATAATCCGGATGCGCTCGGTACCGTCGCATTTGGTCTCGATCCCCAGGCACGCTTAGACATGGGCTAC<br>AACTCCTATGATCAGGTGACTCAGGCCGGCAAAGCCGCCGTTGCACGCTTTTATGGTCGCGCAGCCGACAAGAGCTAC<br>TTCATCGGCTGTTCGGAGGGCGGCCGCGAGGGCATGATGCTGTCCCAGCGCTTTCCATCACATTACGATGGCATTGTG<br>GCGGGCGCACCGGGATATCAGTTGCCGAAGGCCGGAATTAGTGGCGCGTGGACCACCCAGAGCTTAGCGCCCGCCGCC<br>GTTGGCCTGGATGCCCAGGGAGTGCCGCTGATTAATAAGAGCTTTTCTGACGCAGACCTCCATTTACTGTCGCAGGCG<br>ATTCTCGGAACATGCGACGCCTTGGATGGCCTGGCCGACGGCATCGTTGACAACTACCGAGCGTGCCAAGCGGCTTTT<br>GATCCGGCGACTGCAGCCAACCCAGCGAATGGCCAAGCCCTGCAGTGCGTGGGCGCAAAGACAGCCGATTGCTTATCG<br>CCCGTCCAAGTTACGGCGATTAAACGAGCGATGGCCGGTCCGGTAAATAGCGCGGGTACGCCGTTATATAATAGATGG<br>GCCTGGGACGCAGGTATGAGCGGTCTTAGTGGTACCACTTACAATCAGGGTTGGCGCAGCTGGTGGCTGGGATCGTTT<br>AACAGCTCGGCGAATAACGCACAACGTGTATCTGGTTTCTCAGCGCGGAGCTGGCTGGTGGACTTTGCTACCCCGCCG<br>GAGCCGATGCCCATGACCCAAGTCGCCGCCCGTATGATGAAATTTGATTTCGATATCGATCCTCTGAAAATATGGGCT<br>ACTTCGGGCCAATTTACCCAGAGTAGTATGGACTGGCACGGTGCCACTAGCACCGACCTTGCTGCCTTTCGGGACCGC<br>GGCGGTAAAATGATTCTGTATCACGGAATGAGCGATGCCGCATTCTCTGCACTAGATACAGCAGATTATTATGAACGC<br>CTGGGTGCCGCAATGCCGGGCGCCGCGGGCTTTGCTCGTCTGTTCTTGGTTCCGGGAATGAACCATTGCTCCGGGGGT<br>CCAGGTACCGACCGCTTTGATATGCTAACACCGTTAGTTGCATGGGTTGAACGTGGGGAAGCCCCTGACCAAATTAGC<br>GCCTGGAGCGGCACCCCCGGCTACTTTGGTGTGGCCGCCCCGCACTCGACCGTTATGTCCCTATCCGCAGATTGCGCGC<br>TATAAGGGATCAGGCGATATCAATACCGAAGCAAATTTTGCGTGTGCCGCTCCACCGGGTGGTGGTTCTGGTGGTTCT<br>GGTGGTGGTTCTGGTGGTGGTGGTTCTGGTGGTTCTGGTCAGACCAATCCGTATGCGCGCGGCCCCAACCCTACCGCC<br>GCCTCGTTGGAAGCCAGCGCGGGACCCTTTACCGTTCGTAGCTTTACCGTTAGCCGTCCGTCCGGATATGGTGCAGGG<br>ACCGTCTATTACCCAACCAATGCAGGCGGCACCGTTGGCGCGATTGCAATCGTCCCCGGGTACACCGCGCGTCAAAGC<br>AGCATTAAGTGGTGGGGTCCGCGCTTAGCTAGCCATGGCTTTGTGGTTATTACCATCGATACGAACAGCACTCTAGAC<br>CAGCCCAGCAGCCGTAGCTCGCAACAGATGGCCGCGCTTCGTCAAGTTGCGAGCTTGAACGGGACCAGCAGTAGCCCG<br>ATTTACGGAAAGGTCGATACTGCCCGCATGGGTGTGATGGGCTGGTCAATGGGGGGCGGCGGTTCACTTATTAGCGCC<br>GCGAACAACCCGAGTTTAAAAGCAGCGGCACCGCAGGCGCCATGGGACTCTTCAACCAACTTCAGCAGTGTTACCGTG |

| | |
|---|---|
| | CCGACGCTGATTTTCGCGTGCGAGAATGATAGCATTGCACCGGTGAACAGCAGCGCGCTGCCGATTTATGATAGCATG<br>TCCCGCAACGCAAAACAGTTTCTGGAAATTAACGGCGGTAGCCACTCTTGTGCCAACTCTGGGAACAGCAACCAGGCA<br>CTGATCGGAAAAAAAGGGGTTGCATGGATGAAACGATTCATGGATAATGACACCCGTTACTCAACCTTCGCCTGTGAG<br>AATCCCAACAGCACACGCGTGTCGGATTTTCGCACCGCGAACTGTTCCTGA |
| amino acid | MSRDTGDNLDRNQSGNLPMANVMDAYLSRRSVMRGSLGAAIAMIAGTGLTGCVPGDPCAGGGSTPLPLPQQQPPQQEP<br>PPPPVPLASRAACEALKDGNGDMVWPNAATVVEVAAWRDAAPATASAAALPEHCEVSGAIAKRTGIDGYPYEIKFRLR<br>MPAEWNGRFFMEGGSGTNGSLSAATGSIGGGQIASALSRNFATIATDGGHDNAVNDNPDALGTVAFGLDPQARLDMGY<br>NSYDQVTQAGKAAVARFYGRAADKSYFIGCSEGGREGMMLSQRFPSHYDGIVAGAPGYQLPKAGISGAWTTQSLAPAA<br>VGLDAQGVPLINKSFSDADLHLLSQAILGTCDALDGLADGIVDNYRACQAAFDPATAANPANGQALQCVGAKTADCLS<br>PVQVTAIKRAMAGPVNSAGTPLYNRWAWDAGMSGLSGTTYNQGWRSWWLGSFNSSANNAQRVSGFSARSWLVDFATPP<br>EPMPMTQVAARMMKFDFDIDPLKIWATSGQFTQSSMDWHGATSTDLAAFRDRGGKMILYHGMSDAAFSALDTADYYER<br>LGAAMPGAAGFARLFLVPGMNHCSGGPGTDRFDMLTPLVAWVERGEAPDQISAWSGTPGYFGVAARTRPLCPYPQIAR<br>YKGSGDINTEANFACAAPPGGGSGGSGGGSGGGGSGGSGQTNPYARGPNPTAASLEASAGPFTVRSFTVSRPSGYGAG<br>TVYYPTNAGGTVGAIAIVPGYTARQSSIKWWGPRLASHGFVVITIDTNSTLDQPSSRSSQQMAALRQVASLNGTSSSP<br>IYGKVDTARMGVMGWSMGGGGSLISAANNPSLKAAAPQAPWDSSTNFSSVTVPTLIFACENDSIAPVNSSALPIYDSM<br>SRNAKQFLEINGGSHSCANSGNSNQALIGKKGVAWMKRFMDNDTRYSTFACENPNSTRVSDFRTANCS |

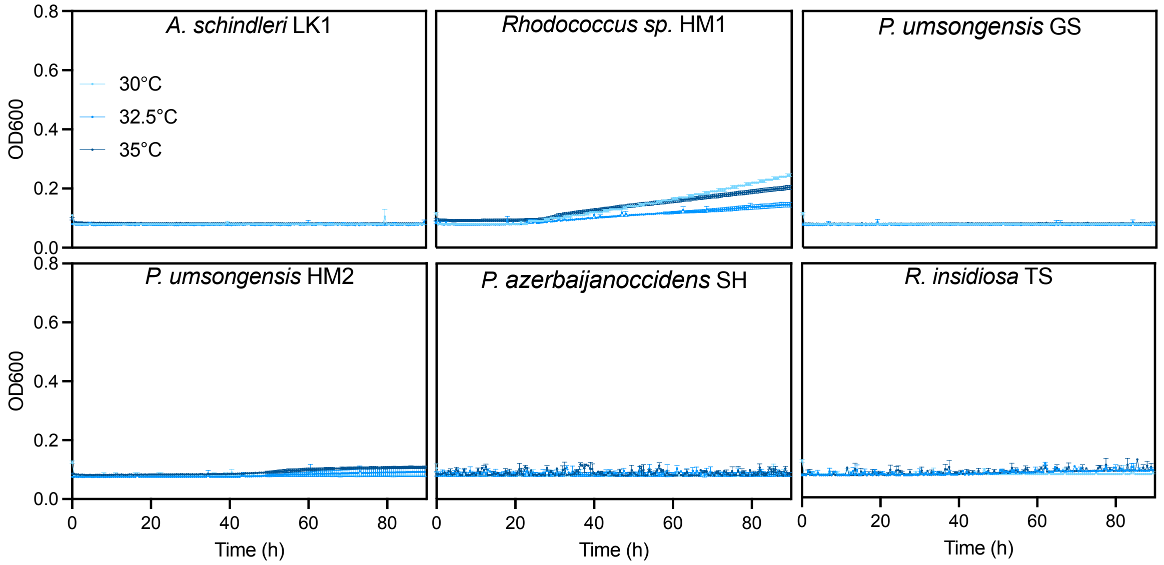


**Fig. S1.** Growth of the environmental isolates using the PET monomer EG as the sole carbon source across a temperature range.

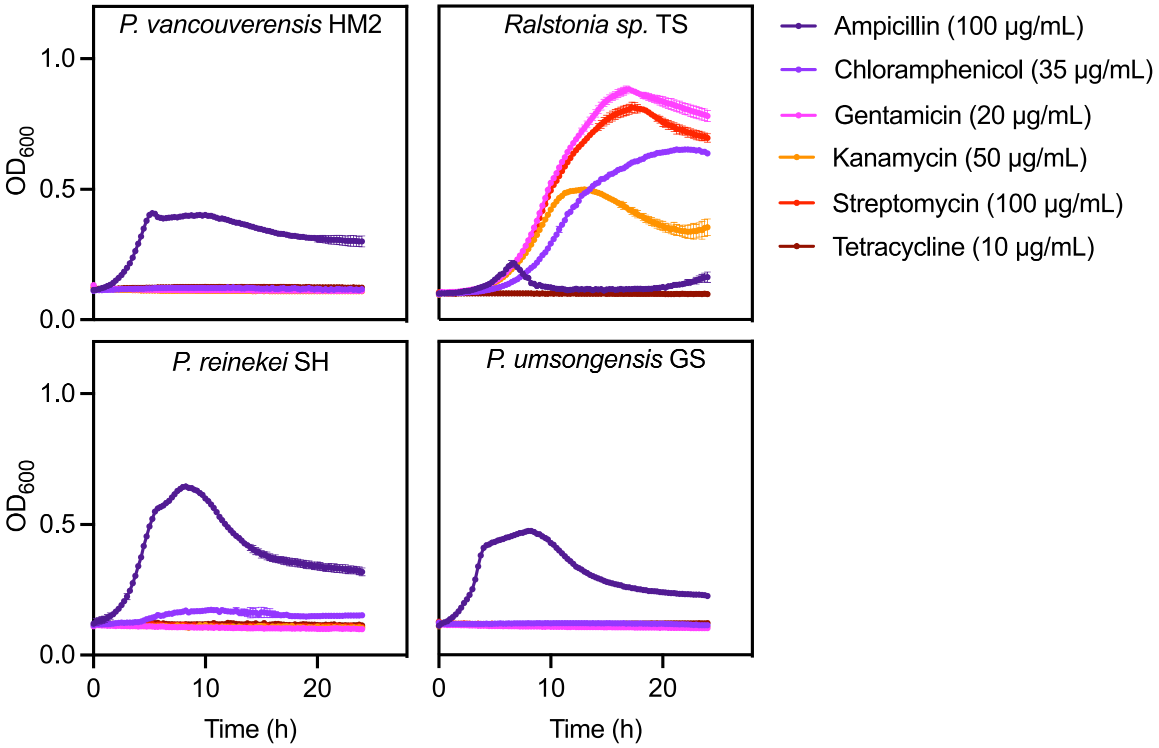


**Fig. S2.** Growth of environmental isolates in the presence of common antibiotics used as selection markers in molecular biology.

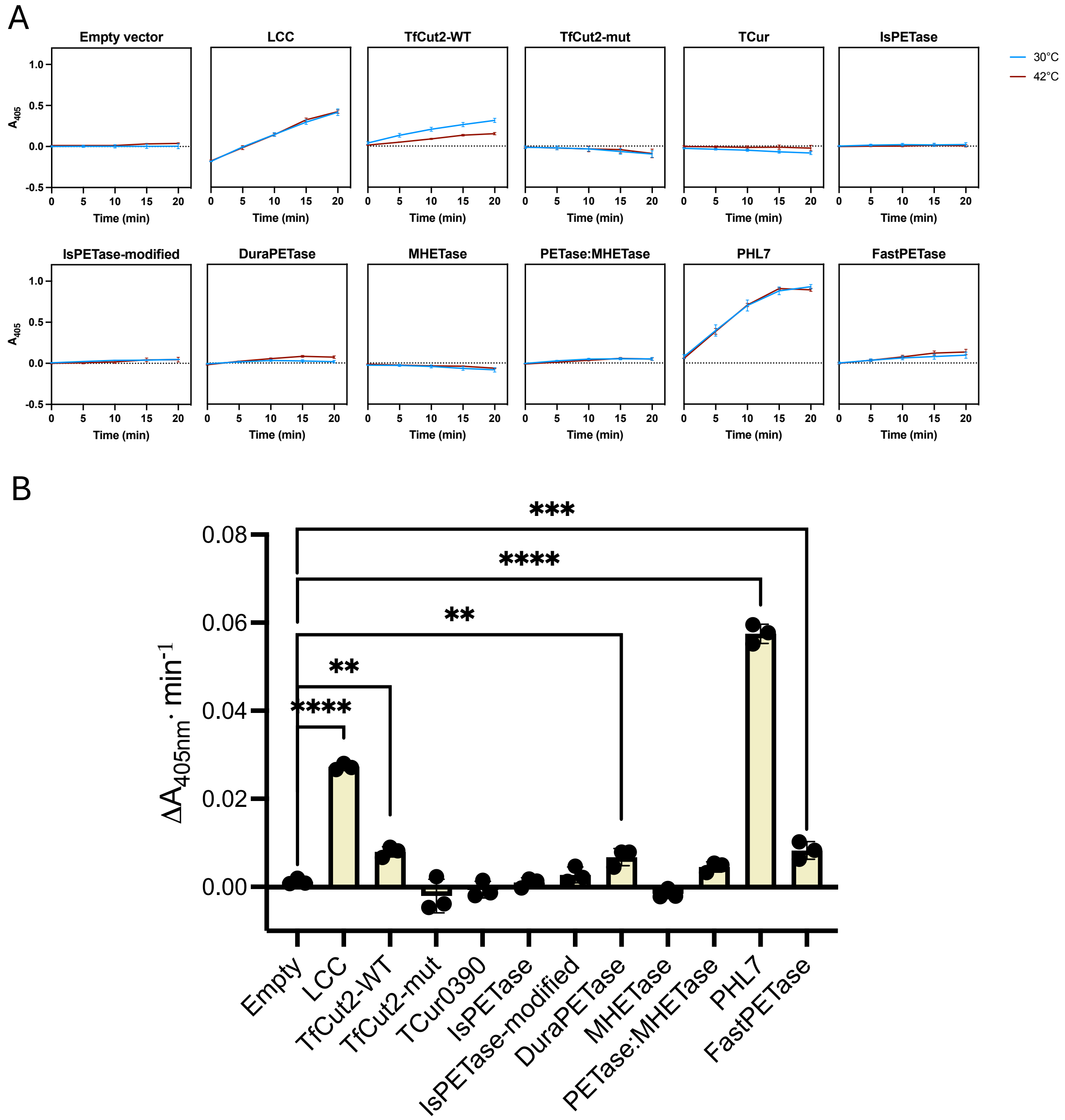


**Fig. S3.** (A) Kinetics of *p*NPB degradation assays at two different temperatures. (B) Activity against pNPB at 42°C of *E. coli* supernatants producing the different enzymes. Error bars indicate the standard deviation of three independent replicates. Within group differences were analysed with an ANOVA followed by a post-hoc Dunnett test. For significance, please check the methods section.

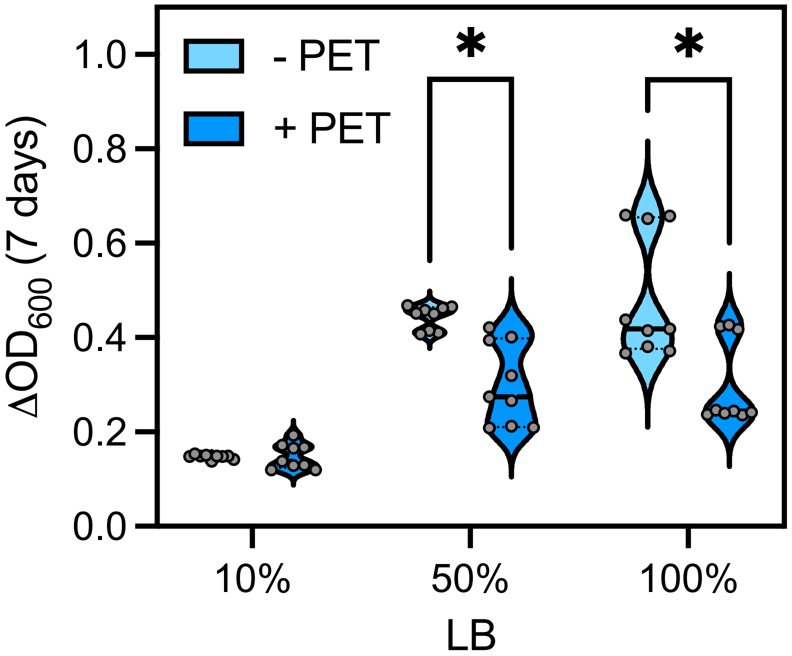


**Fig. S4.** Growth of *P. umsongensis* GS transformed with the empty plasmid pSEVA238 using PET as a growth substrate. Increment in optical density of the strain containing the empty plasmid after 1 week of incubation in the absence (light blue) or presence (dark blue) of activated PET supplemented with different amounts of LB. For significance, please check the methods section.
